# Structure-guided vaccination elicits broadly neutralizing antibodies targeting the HIV-1 silent face

**DOI:** 10.64898/2026.09.30.755568

**Authors:** Joshua J. Carter, Maria B. Palacio, Mark Berlaga, Okikiola Olajide, Joseph J. Noh, Yu Erica Lee, Israel Ramos, Michael R Eso, Jennie Phung, Teresia Chen, Sowmya Meka, Mercedes Guadalupe Garay, Michael S. Seaman, Amelia Escolano, Christopher O. Barnes

## Abstract

An effective HIV-1 vaccine will likely require broadly neutralizing antibodies (bnAbs) targeting multiple complementary epitopes on the Envelope (Env) glycoprotein. Donor-derived bnAbs targeting the conserved silent face (SF) epitope complement established bnAb classes, but the maturation pathways underlying SF bnAb development and their inducibility via vaccination remain poorly understood. Here, we elicit SF-like bnAbs by vaccination and define pathways by which these responses can be guided toward breadth. Structural analysis of the donor-derived bnAb SF10 revealed that broad neutralization depends on a minimal set of heavy chain mutations despite extensive overall somatic hypermutation. We leveraged these insights to engineer Env immunogens that engage an SF10 precursor and selectively favor breadth-associated mutations during sequential immunization in a new immunoglobulin knock-in mouse model. Vaccination reproducibly elicited heterologous neutralizing serum responses and monoclonal antibodies that acquired key SF-like improbable mutations and broad neutralizing activity. Notably, elicited antibodies combined features of distinct human SF bnAbs, demonstrating that vaccination can access alternative mutational solutions rather than precisely recapitulating a donor-derived maturation pathway. These findings establish the silent face as a tractable vaccine target and expand the bnAb specificities available for multi-epitope HIV-1 vaccine strategies.

## INTRODUCTION

Despite advances in HIV-1 prevention and long-acting antiretroviral therapies, achieving durable protection against HIV-1 acquisition remains a major global health challenge, motivating continued efforts to develop an effective preventative vaccine^1,2^. The discovery of broadly neutralizing antibodies (bnAbs) from HIV-1^+^ individuals demonstrated that humoral immunity can achieve remarkably broad HIV-1 neutralization and established bnAbs as a central goal for HIV vaccine development^3,4^. However, despite extensive efforts, reliably inducing bnAb responses in HIV-naïve individuals through vaccination remains a major challenge.

The extensive diversity of HIV-1 Envelope (Env), together with results from passive bnAb administration in high-risk populations, suggests that an effective vaccine will likely require induction of bnAbs targeting distinct, complementary Env epitopes^5^. However, two major barriers have limited progress toward this goal. First, naïve B cell precursors of known bnAbs are often rare and require engineered germline-targeting immunogens for selective activation^6^. Second, many well-characterized bnAbs acquire breadth through prolonged maturation pathways, often accumulating extensive somatic hypermutation (SHM), and in some cases, functionally critical “improbable” mutations^4^. Although sequential immunization strategies have been developed to guide antibody evolution, reliably driving acquisition of key mutations remains an ongoing area of research^7^. These barriers become particularly relevant for multi-epitope vaccination, where complex maturation requirements for individual bnAb lineages could complicate efforts to elicit complementary responses within current bnAb-eliciting regimens^8,9^.

Identifying Env epitopes with more tractable pathways to bnAb elicitation could therefore expand the repertoire of bnAb responses accessible to vaccine development^10^. One such target is the Env silent face (SF), a densely glycosylated surface previously thought to be immunologically inert but now recognized as a target of bnAbs from multiple germline lineages^11–13^. The broadest SF bnAbs identified to date, SF10 and SF12, exhibit neutralization breadth comparable to type I V3-glycan antibodies (∼50-60%), while providing complementary neutralization coverage to this well-studied bnAb class^11,14^. Importantly, prior work suggests that even modest SF-directed responses substantially improve the breadth and potency of polyclonal antibody responses, highlighting their potential value within multi-epitope vaccine strategies^13^. While structural characterization of SF12 provided initial insight into SF recognition, SF12 requires a CDRH2 insertion associated with neutralization, a relatively rare product of antibody maturation that may be difficult to reproducibly elicit by vaccination and reduces its attractiveness as a vaccine target^15^. In contrast, the clonally related bnAb SF10 lacks this requirement while maintaining broad and potent neutralization, albeit with significantly higher degrees of somatic mutations. Whether SF10-like neutralization potency and breadth can be achieved through a more accessible maturation pathway and selectively guided by vaccination remains unknown.

Here, we provide insights into the maturation requirements of SF10-class bnAbs and show that key features associated with broad SF recognition can be selectively driven by vaccination. Structural and functional analyses revealed that SF10 converges on a conserved mode of SF recognition shared with the clonally related bnAb SF12 and that a limited set of SF10 heavy chain mutations is sufficient to support broad neutralization. Using a new immunoglobulin knock-in (Ig KI) mouse model expressing an inferred precursor of SF10 (SF10VHrev), we found that a 3-step sequential immunization strategy with engineered and wild-type (WT) Env immunogens enriched key functional improbable mutations in the SF10 heavy chain and elicited heterologous neutralizing serum and monoclonal antibody responses. Vaccine-elicited antibodies also acquired sequence features shared across distinct SF bnAbs, revealing flexibility in the pathways capable of achieving broad SF recognition. Collectively, our findings define principles governing SF bnAb-like maturation and demonstrate that vaccination can direct antibody responses toward functional features associated with neutralization breadth.

## RESULTS

### SF10 and SF12 converge on a shared mode of silent face recognition

The donor-derived silent face antibody SF10 was previously identified from an elite controller and was part of an expanded clone that utilized VH4-59∗01 and VK3-20∗01 heavy and light chain variable gene segments respectively^15^. SF10 neutralized 51.3% of viruses in a 119-pseudovirus panel representative of major circulating HIV-1 clades with a geometric mean IC50 of 0.1 µg/mL (**Fig. 1A**). While SF10 is clonally related to SF12, it exhibits substantially higher SHM, with amino acid mutation frequencies of 37.1% and 29.0% in the VH and VL regions, respectively, compared to 21.4% VH and 20.4% VL for SF12 (**Fig. 1A**).

**Figure 1.**
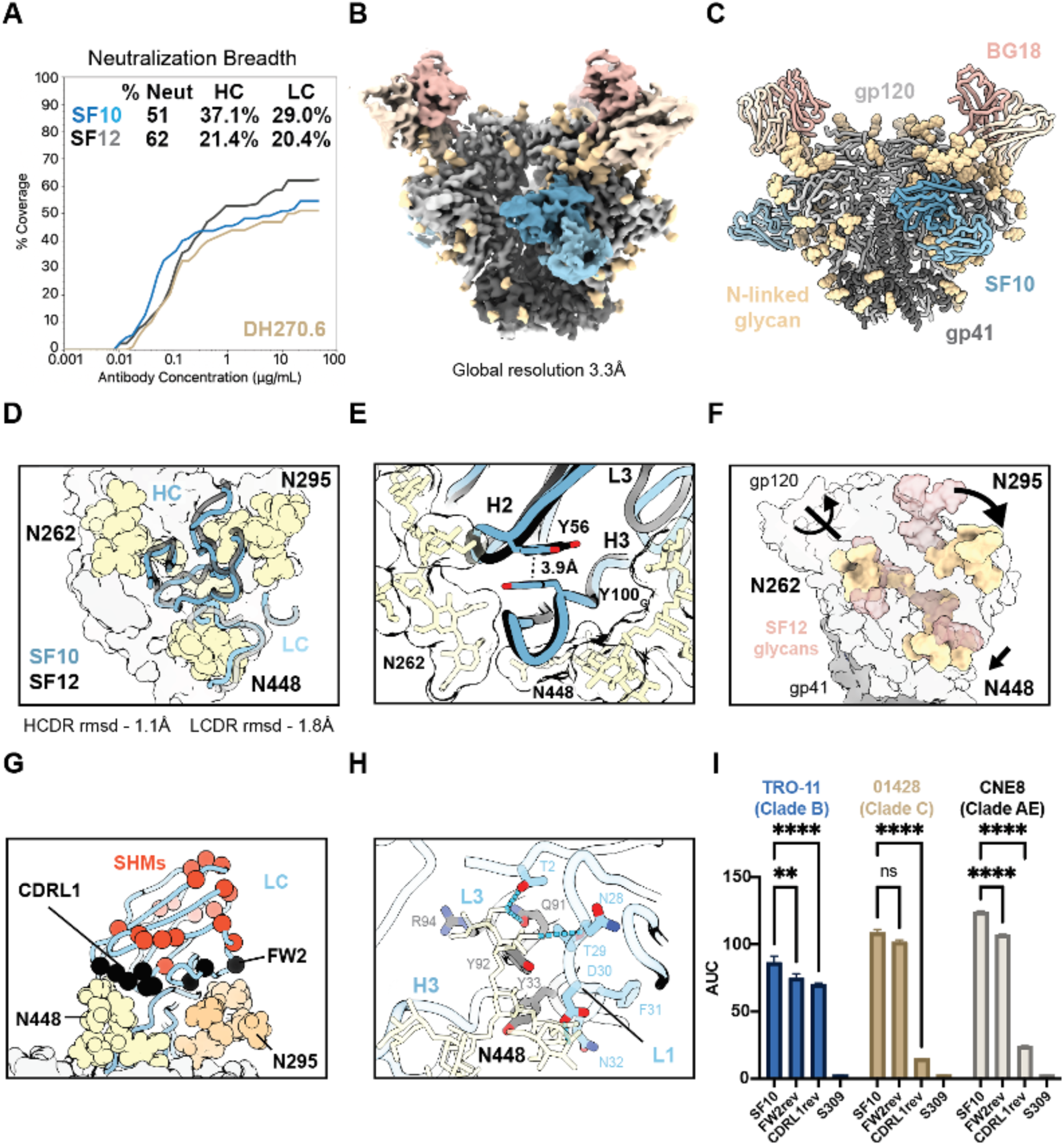
SF10 recognizes the silent face epitope via a CDRH3-dominated binding mode. **A)** Neutralization breadth and potency against a panel of 119 HIV pseudoviruses. **B)** Cryo-EM density map and **C)** atomic model for SF10-BG18-01428 SOSIP structure. **D)** Structural superposition of SF10 (blue) and SF12 (black) showing conservation of CDR loop architecture. Glycans are shown in yellow. **E)** Convergent stabilization of the CDRH3 through pi-stacking interactions in SF10 and SF12. **F)** Comparison of glycan confirmations in SF12 (red) and SF10 (tan) clade B Env-bound cryo-EM structures. **G)** Ribbon representation of SF10 light chain with somatic mutations highlighted as red spheres. Paratope mutations are black spheres. Modeling of the N295 glycan (orange), which is absent in the 01428 strain. **H)** Interactions between CDRL1 and CDRL3 loops with the N448 glycan. Somatic mutations are shown as blue sticks, germline-encoded residues are shown in grey. Potential hydrogen bonds are shown with blue dashed lines. **I)** Binding of SF10 antibody variants to gp120 core proteins measured by ELISA (area under the curve, AUC). Significance was determined from n=3 replicates using 2-way ANOVA and Sidak’s multiple comparisons test. ns p≥0.05, ** p<0.01, **** p<0.0001.

To determine whether the distinct maturation pathways of SF10 and SF12 alter silent face recognition, we determined cryo-EM structures of SF10 bound to clade C (01428) and clade B (TRO-11) Env SOSIP trimers (**Figs. 1B-C, S1A-B**). The resulting structures, solved at 3.3 Å and 3.9 Å global resolution, respectively, revealed a conserved binding mode across divergent Envs. SF10 bound individual gp120 protomers within a cleft between the N262 and N448 glycans, closely mirroring the overall mode of SF12 recognition (**Fig. 1D**)^15^. SF10 recognition was predominantly glycan- and CDRH3-dominated, with glycans contributing 73.4% of total epitope buried surface area (BSA) and the CDRH3 alone contributing over 53.8% of the total paratope BSA and most protein-mediated interactions (74.9% total protein epitope BSA; **Fig. S1C**). Despite sharing only 66% heavy chain (HC) amino acid sequence identity, SF10 and SF12 adopted nearly identical overall CDR orientations and loop architectures (**Fig. 1D**; root mean squared deviation (rmsd) across 37 matched Ca atoms = 1.1Å), suggesting that despite sequence differences, SF10 and SF12 converge on a shared structural solution for broad SF epitope recognition.

This convergence was particularly apparent in stabilization of their 23 amino acid CDRH3 loops. In SF12, a rare tyrosine insertion within CDRH2 is required for neutralization and likely stabilizes the CDRH3 through pi-stacking interactions^15^. In contrast, SF10 achieves a similar outcome through somatic mutation of S56HC to Y56HC, enabling a structurally analogous pi-stacking interaction with Y100G,HC in CDRH3 (**Fig. 1E**). Thus, despite distinct mutational pathways, both antibodies converged on a shared structural solution for stabilizing the CDRH2-CDRH3 interface, suggesting that preservation of this structural interaction may be important for SF recognition.

Despite this conserved antibody architecture, SF10 accommodated substantial variation in the local glycan environment surrounding the silent face epitope. The N262, N295 and N448 glycans adopted distinct conformations relative to the SF12-Env complex (**Fig. 1F**)^15^. Differences in the CDRH1 correlated with altered positioning of the N295 glycan and its interactions with neighboring Env peptide regions, suggesting that somatic mutation differences in the CDRH1 may contribute directly to glycan repositioning while preserving the overall angle of approach and loop architecture. Similarly, the N262 glycan undergoes an approximately 180° concerted rotation of distal mannose branches that is conserved across both clade B and clade C Env-SF10 complexes, indicating that these rearrangements were antibody-driven rather than strain-specific (**Figs. 1F, S1D**). SF10 also maintained a nearly identical binding pose in the clade C 01428 Env SOSIP bound structure despite replacement of the N295 glycan with a glycan at N442 within the epitope footprint (**Fig. S1E**). While this substitution produced subtle shifts in CDRH1 position, it did not disrupt the overall binding mode, suggesting that SF10 can accommodate variation in glycan composition surrounding the silent face with a conserved binding pose.

An important distinction between SF10 and SF12 lies in light chain-mediated glycan recognition (**Fig. 1G**). The SF10 light chain displayed more interactions with the glycan epitope, burying 286Å^2^ and 98Å^2^ total surface area against the N448 and N295 glycans, respectively (compared to 125Å^2^ and 0Å^2^ for SF12), through both germline-and somatic mutation-mediated contacts (**Fig. 1G,H**). To determine whether these interactions contribute to breadth, we reverted SF10 light chain mutations and measured binding to gp120 cores from multiple clades. Reversion of framework region (FWR) 2 mutations contacting the N295 glycan minimally affected binding, whereas reversion of CDRL1 mutations contacting the N448 glycan led to a significant loss of binding (**Fig. 1I**). Single-site reversions identified D30SLC as sufficient to abolish binding to non-clade B strains, consistent with its structural role in stabilizing the N448 glycan (**Figs. 1H, S1F**). Thus, although light chain interactions can contribute to SF10 breadth, their distinct glycan contacts and mutational requirements compared with SF12 indicate greater flexibility in how the light chain accommodates the SF glycan surface.

Collectively, these structural studies revealed that SF10 and SF12 converge on a common mode of recognition despite substantial sequence divergence and variation in glycan composition surrounding the silent face epitope. While their heavy chains adopt conserved structural features associated with broad recognition, their light chains differ in glycan contacts and dependence on somatic mutations, suggesting that antibody breadth may tolerate greater structural plasticity in light chain recognition of the silent face epitope.

#### A limited set of heavy chain mutations underpins SF10 breadth

Given the conserved structural heavy chain elements shared by SF10 and the less mutated SF12, we sought to define the minimal set of mutations required for SF10 neutralization breadth and potency. We first reverted all heavy chain mutations outside the paratope to germline, except for P84, which was conserved across SF lineage members. The resulting paraSF10 variant retained only 17 of the 46 mature heavy chain mutations (10 VH-, 5 D, and 2 J gene-encoded mutations, **Fig. 2A,B**). Because the exact D gene assignment for SF10 is ambiguous between IGHD3-3*01 or IGHD3-9*01, we could not confidently infer germline residues within this region and therefore focused subsequent minimization efforts on VH- and JH-encoded regions. paraSF10 retained binding to clade B (TRO-11) and weak binding to clade C (01428) gp120 cores, but exhibited faster dissociation kinetics and reduced neutralization breadth and potency (**Fig. 2E,F**), indicating that mutations outside the paratope contribute to SF10 function.

**Figure 2.**
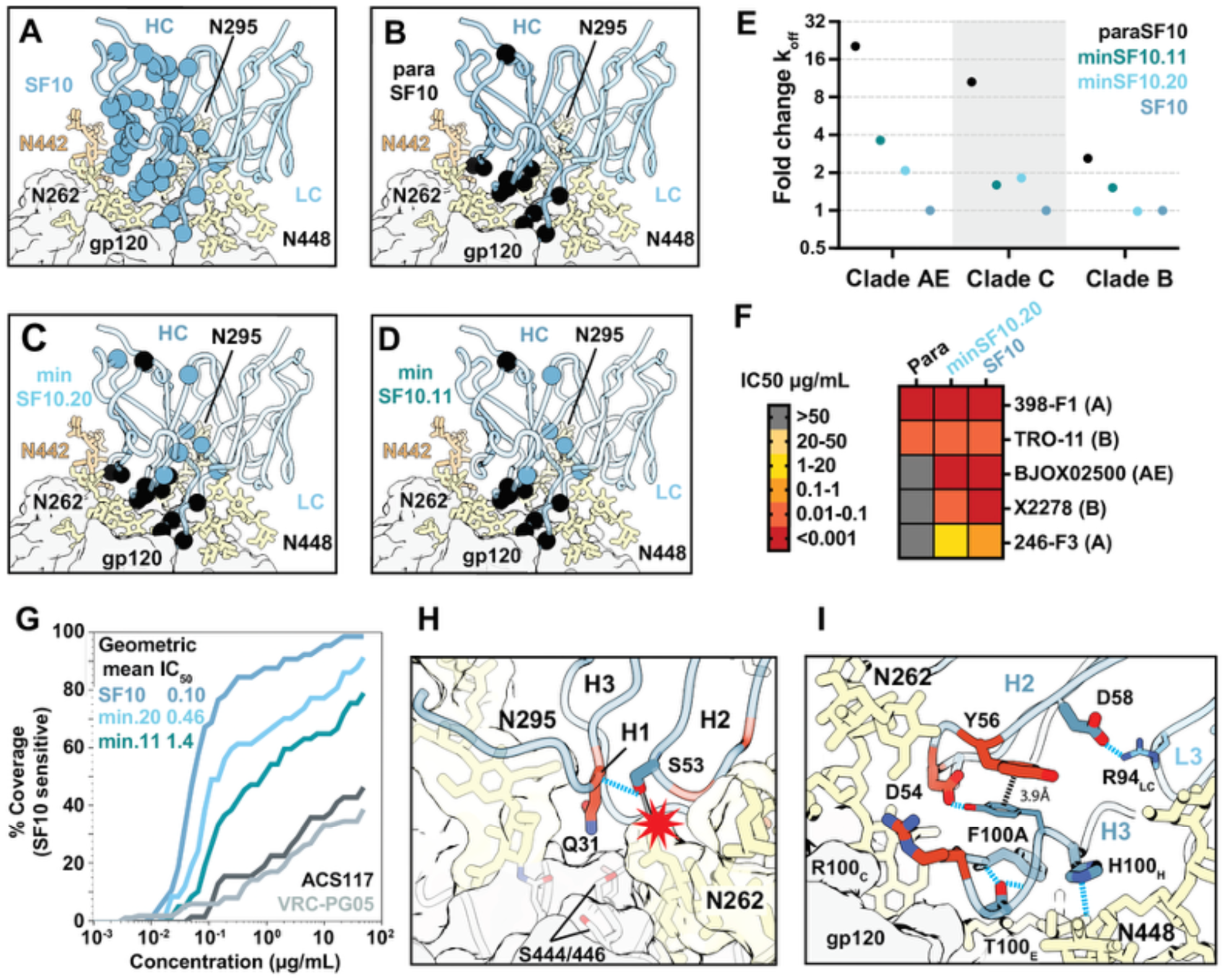
Identification of key heavy chain somatic mutations to confer broad SF10 neutralization. A-D) Ribbon representation of SF10 and minimized variants, with retained somatic mutations shown as spheres. Black spheres denote residues maintained in paraSF10. Color labels correspond to scheme in panel E. **E)** Fold change in dissociation rates (k_off_) of heavy chain variants binding gp120 cores from multiple HIV-1 clades relative to mature SF10. **F)** Neutralization heat map comparing para, minSF10.20 and mature SF10 against the five SF10-sensitive viruses within the 12-strain global panel. **G)** Neutralization breadth and potency of minimized SF10 variants against the 65 SF10-sensitive viruses from the 119 HIV pseudovirus panel compared to mature silent face bnAbs SF10, ACS117, and VRC-PG05. Reported IC_50_ values represent geometric mean IC50 values for all 65 strains. **H,I)** Structural basis of key breadth-associated HC somatic mutations. “Improbable” mutations predicted by ARMADILLO are shown in red. Blue dashed lines denote potential hydrogen bonds. Red star denotes a predicted steric clash of the germline Y53 residue and the N262 glycan.

To identify functionally important non-paratope mutations, we generated a single-chain variable fragment (scFv) yeast display library in which the remaining 29 reverted heavy chain positions varied between germline and mature residues while maintaining the mature light chain (theoretical library size 5.36 x 10^8^). Parallel selections against gp120 cores (gp120c) from clade AE (CNE55), clade B (B41), and clade C (01428), followed by deep sequencing, identified a limited subset of enriched mutations shared across libraries sorted with different gp120c. Stepwise incorporation of these mutations into paraSF10 progressively restored binding and neutralization breadth, yielding minSF10.20 (containing 20 non-D gene mutations), which showed broad neutralization activity against clade B, C, AE, and A viruses comparable to mature SF10 (**Fig. 2C,E,F**).

To further refine the minimal requirements for breadth, we next individually reverted the remaining VH- and JH-encoded mutations in minSF10.20 and assessed binding across multiple gp120 cores (**Fig. S2A**). This analysis identified seven VH mutations, corresponding to only 11% amino acid divergence from germline VH, that were sufficient to preserve broad neutralizing activity. This minimized antibody construct, minSF10.11, maintained high-affinity binding across all tested gp120s (**Fig. 2D,E; Table S1**). When evaluated against 65 SF10-sensitive viruses, minSF10.20 and minSF10.11 neutralized 93% and 78%, respectively, corresponding to 47.2% and 39.5% breadth on the global panel, with geometric mean IC50 values comparable to the mature SF10 (**Fig. 2G**). Both minimized constructs also outperformed previously described silent face bnAbs VRC-PG20 and ACS117^12,13^. Thus, the extensive SHM accumulated by mature SF10 substantially exceeds the mutational requirements necessary to reproduce much of its neutralization breadth and potency.

Mapping functionally important mutations from minSF10.11 onto the SF10 structure suggested potential mechanistic roles for individual substitutions in supporting antibody breadth. The Y53S CDHR2 mutation prevents a predicted steric clash with Env glycans, while additional CDRH2 and FWR3 mutations likely stabilize interactions between the CDRH1, CDRH3, and CDRL3 loops (**Fig. 2H,I**). Similarly, the S31QHC mutation is positioned to enhance epitope engagement by filling a pocket formed by the N295 glycan and adjacent Env residues (**Fig. 2H**). Three of these mutations (S31QHC, S54DHC, and S56YHC) are predicted by ARMADILLO to be improbable (<1% frequency of occurrence in 10,000 simulations of equivalently mutated antibodies in the absence of selection; **Fig. 2H,I**)^16^. These sites are also mutated to identical or similarly improbable residues in SF12, consistent with shared functional constraints despite the distinct maturation pathways of these antibodies. In contrast, a minimally mutated light chain containing only four probable mutations at the N448 binding site maintained neutralization across multiple clades, suggesting a more attainable maturation pathway for the light chain (**Fig. S2D**). Together, these analyses define a limited set of heavy chain somatic mutations that underpin SF10 breadth and potency, including improbable mutations that may present barriers to vaccine-mediated maturation. Conversely, the minimal light chain mutations required for broad activity were all probable, motivating our subsequent focus on heavy chain maturation *in vivo*.

#### Structure-guided immunogen design enables engagement of the SF10VHrev precursor

The concentration of multiple functionally important and improbable mutations within the SF10 heavy chain prompted us to generate an *in vivo* model to test whether potential regimens could select these features through vaccination. We designed a partially reverted SF10 heavy chain precursor (SF10VHrev) in which the entire VH and FWRH4 were reverted to their inferred germline sequences while the CDRH3 and mature light chain were retained (**Figs. 3A, S3A**). As longitudinal lineage sequences were unavailable, the CDRH3 was preserved due to ambiguity in D gene assignment and the fact that the remaining J gene mutations were predicted by ARMADIILLO to arise with high probability through SHM^16^. This design therefore minimized heavy chain mutations while preserving sequence features that could not be confidently reverted or were unlikely to represent major maturation barriers. SF10VHrev mAb lacked neutralizing activity against the 12-strain global virus panel and bound clade B and clade C gp120 cores weakly (**Fig. S3B**), indicating that native Env was unlikely to efficiently engage this precursor. We therefore sought to engineer an immunogen with enhanced affinity for SF10VHrev.

**Figure 3.**
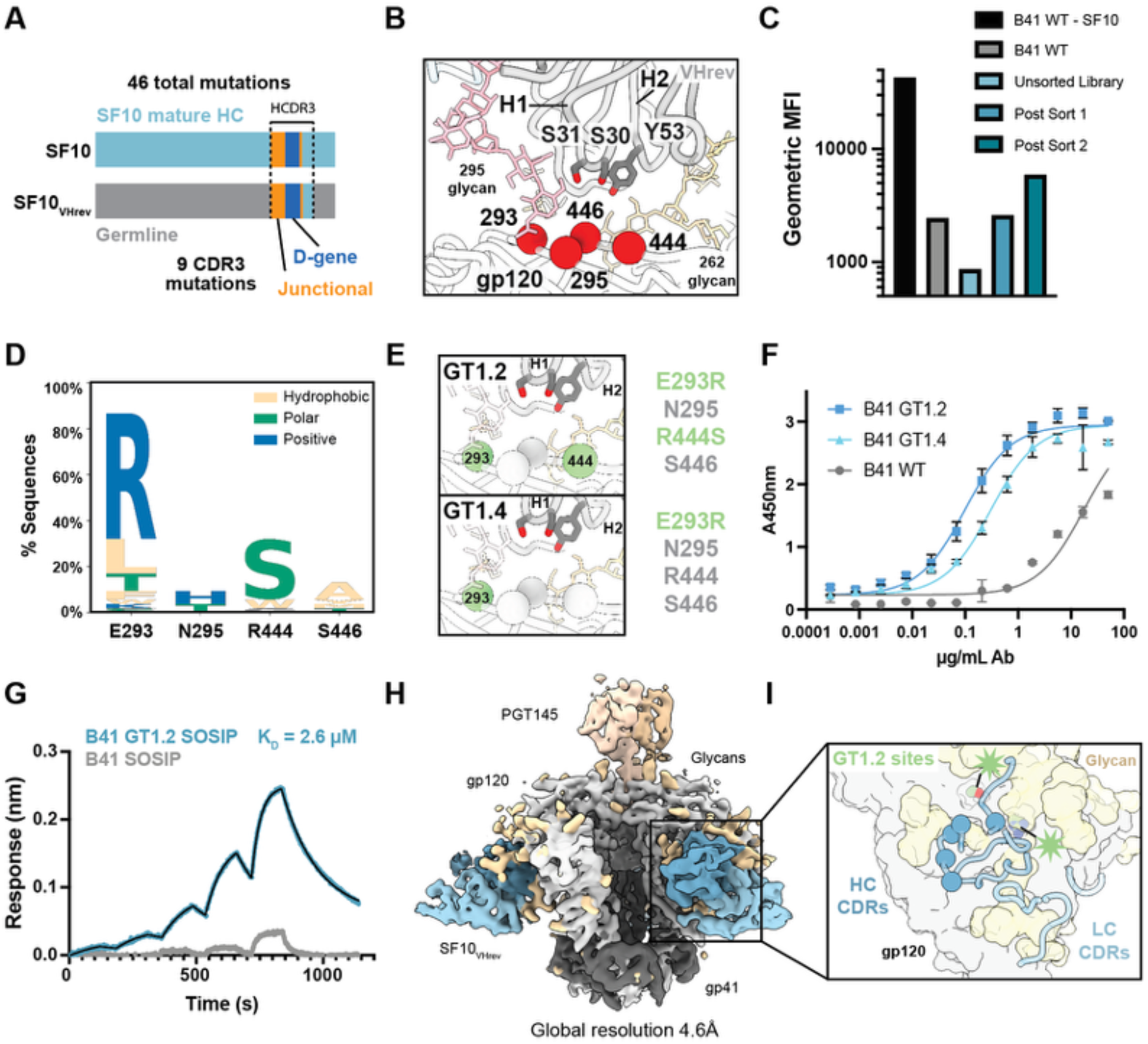
Engineered and native Env immunogens engage SF10VHrev precursors. **A)** Schematic showing the regions outside the CDRH3 reverted in the SF10_VHrev_ antibody model. Detailed sequence information on SF10_VHrev_ is shown in Supplemental Fig 3A. **B)** Structural location of sites chosen for randomization in yeast display library. Sticks highlight relevant paratope residues that differ between SF10_VHrev_ and SF10. **C)** Geometric median fluorescence intensity for parental strain and libraries over the directed evolution campaign. **D)** Sequence logo plot showing selected sequences at the four randomized sites. The germline residue is left blank for clarity and colors are defined as shown. **E)** Locations of selected sites in the two highest affinity selected clones by ELISA, B41 GT1.2 and GT1.4. Sequences at the randomized sites are shown adjacent. Mutated sites are shown in green, sites that retain the wildtype residue are grey. **F)** ELISA of SF10_VHrev_ IgG binding to B41 GT1.2, GT1.4, and WT gp120c. Monomeric affinities of the gp120c immunogens were determined via BLI and are available in Supplemental Figure 3. B41 GT1.2 was selected for further characterization and development. **G)** Single cycle kinetics of B41 GT1.2 or WT Env trimers binding to SF10_VHrev_ Fab. Fitted monomeric KD is reported for B41 GT1.2 SOSIP. **H)** Structure of SF10_VHrev_ binding to B41 GT1.2 SOSIP. The structure was resolved to 4.6Å and a crystal structure of SF10_VHrev_ was docked in to facilitate modelling of interactions. **I)** The location of SF10_VHrev_ CDRs is shown with the identified key heavy chain mutation sites highlighted as blue spheres. Sites mutated in GT1.2 are highlighted with green stars for clarity.

Guided by the SF10-Env structures and the previously published SF12-B41 structure, we generated a yeast surface display library of B41 gp120c variants containing targeted saturation mutagenesis within the SF epitope (**Fig. 3B**). Selection for enhanced SF10VHrev binding enriched substitution at Env positions 293 and 444 (**Fig. 3C,D**). Characterization of the two most enriched variants, B41 GT1.2 and GT1.4, highlighted a shared E293R mutation in Env that is positioned near the CDRH1 and -H3 loops (**Fig. 3E**), with the lead immunogen, B41 GT1.2, containing both E293R and R444S. *In vitro* characterization of these variants demonstrated improved affinity relative to B41 WT gp120c proteins, with B41 GT1.2 showing increased binding to SF10VHrev by more than 100-fold by ELISA and micromolar monomeric affinity by BLI (8.6 µM; **Figs. 3F, S3C,D**). Introduction of the two GT1.2 mutations into a B41 MD39 SOSIP background similarly enhanced SF10VHrev binding to trimeric Env (monomeric KD 2.6µM; **Fig. 3G**).

To understand the structural basis for enhanced precursor engagement, we determined a 4.5 Å cryo-EM structure of SF10VHrev bound to B41 GT1.2 SOSIP and solved the SF10VHrev Fab crystal structure to 2.35 Å resolution to facilitate modeling at the Fab-antigen interface (**Fig. 3H,I**; **Tables S2,3**). While the resolution of our cryo-EM structure precluded assignment of side-chain level interactions, the engineered R293 substitution is positioned to bridge the N295 glycan and germline CDRH1 of SF10VHrev, providing a plausible structural explanation for enhanced binding (**Fig. S3E**). The structure also revealed an extended and ordered glycan network surrounding the SF epitope involving the N295, N332, N392, and N413 glycans, suggesting that both local protein contacts and glycan organization likely contribute to enhanced engagement (**Fig. S3F-H**).

As monomeric B41 GT1.2 affinity for SF10VHrev was still relatively low (8.6 µM), potentially reducing effective B cell activation and competition in germinal centers^17^, we evaluated whether multivalent immunogen display could enhance precursor recognition (**Fig. S3I**). B41 GT1.2 gp120c displayed on mi3 or ferritin nanoparticles exhibited robust avidity-enhanced SF10VHrev binding with negligible dissociation (**Fig. S3I**). In contrast, dense display of B41 GT1.2 SOSIP on mi3 nanoparticles impaired SF10VHrev IgG binding, but not Fab, consistent with steric constraints imposed by a lower angle of approach. SOSIP trimer display on the lower valency ferritin nanoparticle restored robust binding to SF10VHrev IgG, albeit with a faster off rate than B41 GT1.2 gp120c displayed on mi3 (**Fig. S3J**). Thus, these findings demonstrate that both affinity, antigen-display and glycan context influence efficient engagement of the SF10VHrev precursor.

Having established immunogens capable of engaging SF10VHrev, we generated immunoglobulin knock-in (Ig KI) mice expressing this precursor to evaluate SF10 heavy chain maturation *in vivo* using a newly developed method (submitted; **Fig. S4A**). Naive SF10VHrev Ig KI mice displayed normal B cell development and efficient expression of the knocked-in B-cell receptor, with 97.1% of peripheral blood mature B cells binding the B41 GT1.2 gp120c probe (**Fig. S4B-D**). Thus, SF10VHrev Ig KI mice provide a functional *in vivo* model for testing whether immunization can select heavy chain mutations associated with SF10 breadth.

#### A 3-step sequential immunization protocol induces heterologous neutralization in SF10VHrev mice

To determine whether sequential immunization could drive SF10VHrev responses toward breadth, we primed SF10VHrev IgKI mice with our engineered B41 GT1.2 gp120c-conjugated mi3 nanoparticle followed by native-like Env DS-SOSIP-conjugated ferritin boosts selected to favor acquisition of breadth associated mutations (**Fig. 4A**). Whereas GT1.2 efficiently engaged SF10VHrev, wild-type Env trimers bound the precursor weakly but showed increased affinity for antibody variants containing key improbable VH mutations (**Fig. 4B**). We therefore used these low-affinity native Envs as selective affinity gates, reasoning that B cells acquiring mutations that improved recognition to wild-type Env boost immunogens would be preferentially selected and drive continued affinity maturation. Since no single Env tested displayed increased affinity for all key mutations, we compared two boosting strategies: Group 1 (G1) mice received the autologous clade B B41 trimer, which showed increased affinity for the S31QHC mutation, whereas Group 2 (G2) received the heterologous clade C 01428 trimer, which preferentially showed increased affinity for the improbable S54DHC mutation (**Fig. 4A,B**). 01428 also presents a distinct SF glycan environment, lacking the N295 glycan and containing the N442 glycan, providing an additional selective pressure that we hypothesized could reduce dependance on N295 glycan-mediated interactions with the CDRH1 and favor selection of mutations with broader SF recognition. To further promote acquisition of breadth-associated mutations, both groups received a second boost with the clade AE CNE8 DS-SOSIP-ferritin nanoparticles two weeks after the first boost (w4) (**Fig. 4A**).

**Figure 4.**
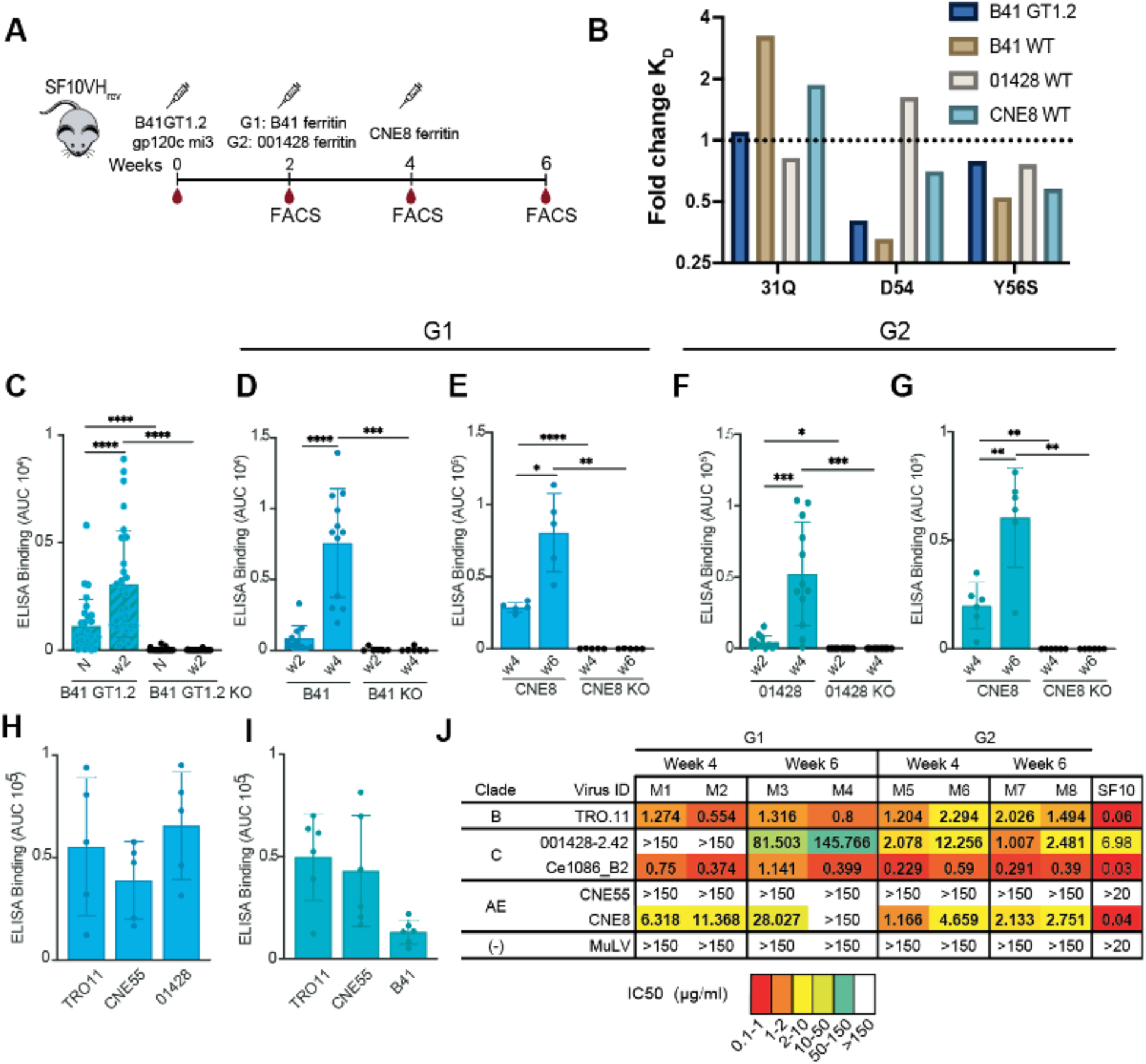
A prime-boost regimen rapidly elicits SF-specific heterologous neutralizing sera responses in a SF10_VHrev_ knock-in mouse model. **A)** Sequential immunization protocol. All mice received a prime immunization with B41GT1.2 gp120 core on mi3 nanoparticles (B41GT1.2 gp120c mi3) and were subsequently divided into two groups as indicated. **B)** Fold change K_D_ for minSF10.11 with key improbable mutation reversions to germline. Values greater than 1 imply increased affinity for a particular mutation over germline by each immunogen strain. **C)** ELISA results showing the binding of the serum from naïve (N) SF10_VHrev_ mice and serum collected two weeks after the binding of prime (w2) to B41GT1.2 gp120c (B41GT1.2) and a knockout version that abrogates the silent face epitope (B41GT1.2 KO). Each dot corresponds to a mouse. n= 29 mice primed with B41 GT1.2 gp120 core. **D,F**) ELISA results showing binding of serum from mice in G1 and G2 respectively collected after the prime (w2) and after the first boost (w4) to the gp120c corresponding to the respective boosting immunogen B41 (G1) or 01428 (G2) and to their corresponding silent face KO protein (B41 KO, 001428 KO) **(E,G)** ELISA results showing binding of serum from mice in G1 and G2 respectively collected after the first boost (w4) and after the second boost (w6) to the boosting immunogen CNE8 gp120c and to the corresponding silent face KO protein (CNE8 KO). **(H,I)** ELISA results showing binding of serum from mice in G1 and G2 respectively to the indicated heterologous Envs. **J)** Results of TZM-bl neutralization assays using purified serum IgGs from the sequentially immunized mice in G1 and G2 against a panel of five tier 2 HIV-1 pseudoviruses. Statistical analysis was performed using a two-tailed Student’s t-test presented as mean with SD. Only significant P values are shown as * p< 0.05, **p< 0.005, ***p< 0.001, ****p<0.0001.

Priming with B41 GT1.2 gp120c-mi3 nanoparticles in SMNP adjuvant elicited robust SF-specific responses in all SF10VHrev mice, with strong serum binding to the priming immunogen but not the B41 GT1.2 gp120c N448K construct, a mutant protein with a knocked-out SF epitope (**Fig. 4C**). Following the prime, mouse sera showed low but detectable binding to the autologous B41 WT gp120c, with reactivity also observed against the heterologous clade C 01428 gp120c, suggesting early recognition of native SF epitopes. Boosting with either B41 (G1) or 01428 (G2) further increased responses to native Env, while maintaining SF specificity (**Fig. 4D,F**). Following the second boost with CNE8, sera from both groups exhibited significantly increased binding and cross-reactivity with multiple heterologous native Env trimers (**Fig. 4E,G-I**).

Importantly, heterologous neutralization was observed after the first boost. By week 4, purified serum IgG from both groups neutralized viruses from clades B, C, and AE, including TRO-11, Ce1086, and CNE8, respectively (**Fig. 4J**). G2 animals additionally neutralized the autologous 01428 strain at w4 and w6, whereas weaker 01428 neutralization emerged in G1 animals only after the heterologous CNE8 boost at w6. Neither group neutralized the clade AE CNE55 strain, consistent with the known specificity profile of SF10. Thus, these findings demonstrated that a rationally selected sequential immunization protocol using native Env affinity gates can rapidly induce SF-specific serum responses with heterologous neutralization across multiple HIV clades, suggesting that a SF10-like precursor can be driven toward functional breadth *in vivo*.

#### Sequential immunization selects for key improbable SF10-like somatic mutations

To determine whether vaccine-stimulated SF10VHrev B cells acquired SHMs identified as critical for SF10 function, we isolated single SF-specific germinal center (GC) B cells from the draining lymph nodes of sequentially immunized SF10VHrev Ig KI mice two weeks after the prime, first (w4), and second (w6) boosts. SF-specific GC B cells were isolated using antigen probes containing intact or knocked-out silent face epitopes (**Fig. 5A**). IgH sequencing revealed substantial somatic diversification at the nucleotide (nt) and amino acid (AA) level after only a single boost, with both mutation frequencies increasing further after the second boost (**Fig. 5B,C**). Mice receiving the heterologous 01428 boost (G2) initially accumulated more mutations than B41-boosted mice (G1), although both groups reached similar mutation frequencies following the final CNE8 boost (**Fig. 5B,C**). Importantly, this diversification was accompanied by progressive enrichment of AA mutations matching those observed in mature SF10 in both groups, with G2 mice acquiring slightly more SF10-like mutations at w4 that persisted through w6 (**Fig. 5D**).

**Figure 5.**
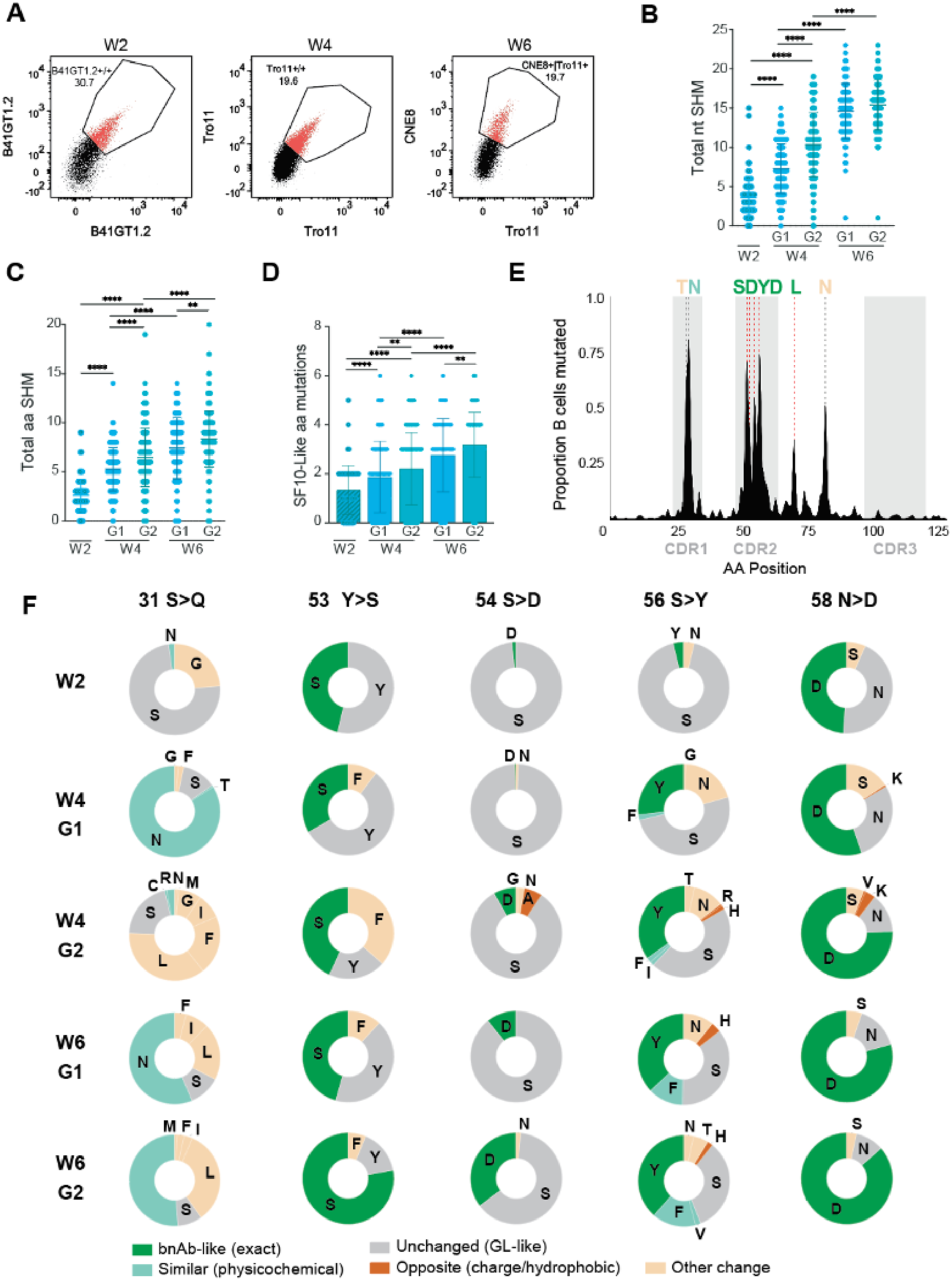
Sequential immunization selects for key functional somatic mutations in SF10_VHrev_ mice. **A)** Representative flow plots showing B41GT1.2^+/+^ (left), Tro11^+/+^ (middle) and Tro11^+^/CNE8^+^ (right) GC B-cells isolated from the immunized mice at w2, w4 and w6 respectively. B cells were pre-gated as B220^+^, Dump^-^; GL7^+^, CD95^+^; Tro11KO^-^. **B,C)** Total number of nucleotide and amino acid somatic hypermutations (SHM) in the heavy chains of single B cells isolated from the sequentially immunized mice in G1 (blue) and G2 (green). Each dot represents one sequence. **D)** Progressive enrichment of SF10-like amino acid SHM in sorted B cells. **E)** Frequency plot of SHM along SF10_VHrev_ heavy chain primary sequence. Predominant amino acids at highly mutated positions are noted above the plot using the color scheme shown in panel F. **F)** Donut plots represent the type of amino acid substitutions observed at key positions for SF10 neutralization and binding at w2, w4 and w6 in mice from G1 and G2. Statistical analysis for B-D was performed using a two-tailed Student’s t-test and presented as mean with SD. Only significant P values are shown as * p< 0.05, **p< 0.01, ***p< 0.001, ****p<0.0001.

We next examined whether vaccination preferentially selected mutations at positions previously identified as important for SF10 function. Mutations accumulated recurrently within CDRH1, CDRH2, and FWR3, with five of the eight most frequently mutated positions corresponding to residues essential in minSF10.11 (**Fig. 5E**). Across several key sites identified in our minimization campaign, including positions 53, 56, and 58, the mature SF10 residue or chemically similar substitutions represented the most abundant mutation observed, with positions 53 and 58 showing enrichment two weeks after the prime (**Fig. 5F**). Notably, position 54 emerged as one of the most frequently mutated residues despite lacking a canonical activation-induced deaminase (AID) hotspot motif, with substitutions converging almost exclusively on the improbable S54D mutation associated with SF10 breadth (**Fig. 5F**). In contrast, the breadth-associated S31Q mutation was not observed, with most antibodies instead acquiring alternative asparagine and leucine amino acids at this position. Interestingly, we did not observe additional diversification within the mature light chain genes of the SF10VHrev Ig KI mice, consistent with our earlier findings that heavy chain maturation represents the dominant determinant of SF breadth acquisition (data not shown). Collectively, these findings demonstrated that sequential immunization did not simply drive extensive somatic mutation, but preferentially selected mutations at positions independently identified as important for broad SF10 function, including improbable SF10-like substitutions. At the same time, selection of alternative residues at position 31 suggested that vaccination could access mutational solutions distinct from the mature SF10 pathway.

### SF10-like mAbs isolated from sequentially immunized SF10VHrev Ig KI mice broadly neutralize HIV-1

To define the functional activity of vaccine-elicited monoclonal antibodies (mAbs), we expressed 30 mAbs comprised of representative immunoglobulin heavy chains isolated at w4 and w6 from G1 and G2 mice paired with the mature SF10 LC. Nearly all mAbs (26/30) bound CNE8 gp120c, with higher affinity among w4 G2 mAbs compared to the w4 G1 mAbs (geometric mean *KD* 105 nM vs 6.08 µM across 8 mAbs from each group, Mann-Whitney p=0.0204; **Fig. S6A**). We selected twelve mAbs spanning distinct expanded clones, binding affinities, and SF10-like mutation profiles for neutralization analysis. All 12 selected mAbs exhibited heterologous neutralization on a multi-clade five-virus panel, demonstrating that the SF10-like maturation observed following immunization translated into broad antiviral activity (**Fig. 6A**). One elicited antibody, 14.1, neutralized more viruses than mature SF10 on this panel despite containing only nine VH mutations and three of the five key SF10 VH mutations. At the remaining two positions, mAb 14.1 acquired S31L, matching the amino acid in mature SF12, and S56F, a physiochemically equivalent substitution to SF10 S56Y. Thus, 14.1 did not precisely recapitulate SF10 maturation but instead combined features of distinct human SF bnAbs while achieving enhanced neutralization breadth on the tested panel.

**Figure 6.**
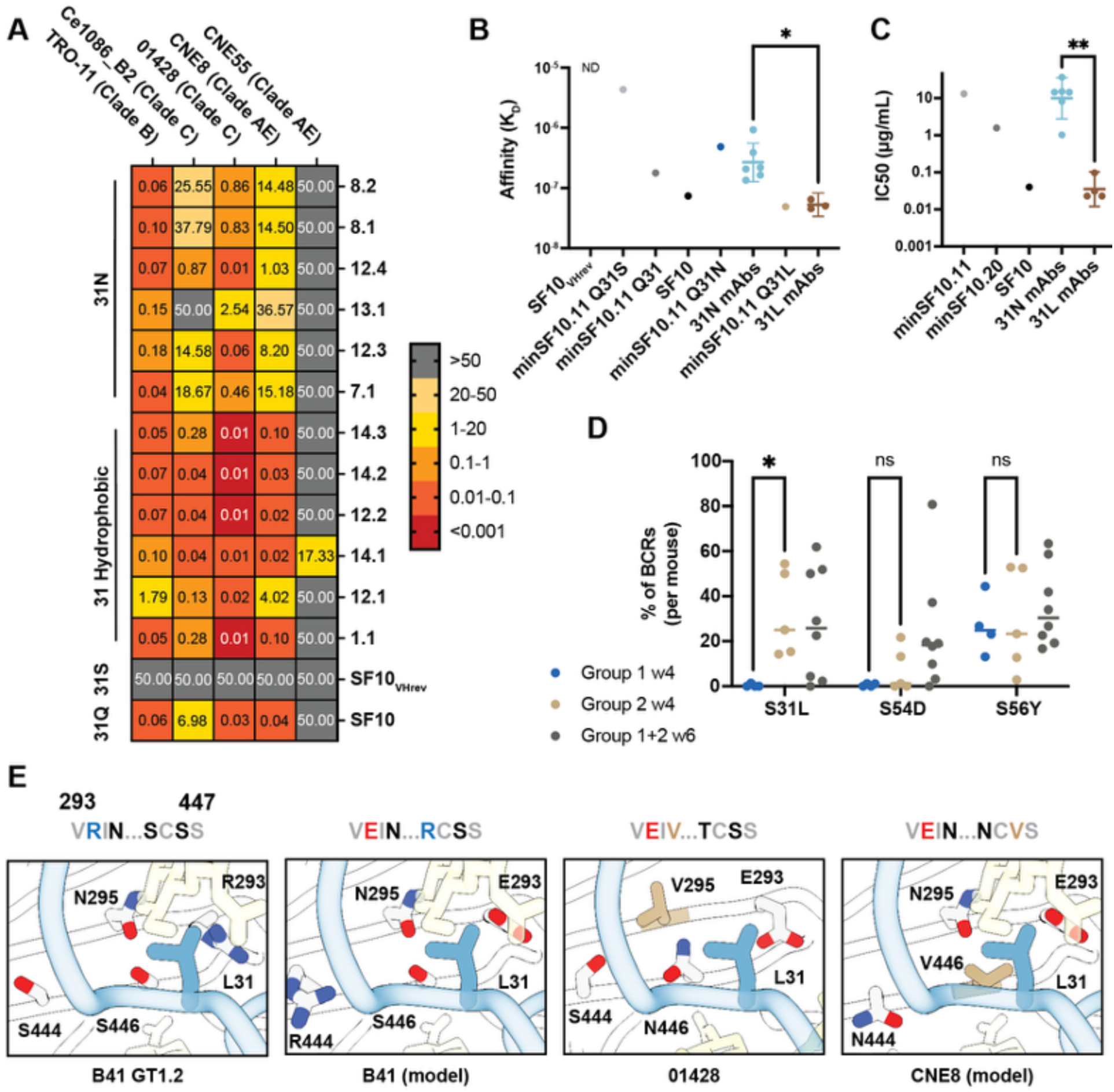
Vaccine-elicited antibodies achieve broad HIV-1 neutralization through alternative SF10 maturation pathways. **A)** Neutralization breadth and potency of representative vaccine-elicited monoclonal antibodies against a multiclade 5-strain panel. Bars denote amino acid character at position 31. **B)** Binding affinities of antibody variants at position 31 to CNE8 gp120c measured by BLI. **C)** Neutralization of CNE8 pseudovirus by mAbs containing alternative amino acid substitutes at position 31. **D)** Frequencies of amino acid mutations reported at position 31 across individual immunized mice. Statistical significance for panels B-D was determined using the Wilcoxon rank sum test with Bonferroni correction for multiple hypothesis testing for panel D. * p<0.05, ** p<0.01. **E)** Structural environment of L31 mutation in immunogen strains, with accompanying sequences adjacent. Hydrophobic amino acids are colored tan. The position of L31 was modeled onto the SF12-B41 structure (PDB ID: 6OKP). CNE8 amino acids at relevant sites were modeled onto the B41 GT1.2 structure for comparison.

We next investigated whether the alternative S31 substitutions selected by vaccination contributed to antibody function. S31L-containing mAbs exhibited more potent neutralization than S31N-containing mAbs (Mann Whitney p=0.0095; **Fig. 6A,C**), while other hydrophobic substitutions at this position, including S31F and S31I, showed similar neutralization profiles to S31L (**Fig. 6A**). To confirm that variation at position 31 was responsible for the observed differences, we introduced S31N or S31L in the minSF10.11 background and evaluated their binding to CNE8 gp120c via BLI (**Fig. 6B**). Both S31N and S31L mutations resulted in improved affinity for CNE8 gp120c relative to the germline S31, with the greater affinity of the S31L-containing construct paralleling the increased neutralization potency of elicited S31L mAbs (**Fig. 6B,C**). Collectively, these results demonstrate that variation at position 31 is a key determinant of SF antibody function and reveal that selection for hydrophobic amino acids at position 31, especially S31L, drives increased breadth and potency.

Finally, given the importance of variation at position 31, we asked whether immunogen choice contributed to selection at this and other key improbable mutation sites. Both 01428 and CNE8 preferentially bound the S31L variant over S31N (**Fig. S6B**), and S31L subsequently comprised up to 54.4% and 61.9% of the total antigen-sorted BCR sequences following 01428 and CNE8 boosts, respectively (Fig. 6D). Consistent with its affinity profile, 01428 drove significantly greater S31L enrichment compared to B41-boosted mice (**Fig. 6D**; Mann-Whitney rank sum p=0.0476 after Bonferroni correction) and also enriched S54D at much higher rates than expected in several mice. Structural analysis suggests that this mutational selectivity may be driven by a more hydrophobic local epitope environment at position 31 in 01428 and CNE8 compared to B41, although this remains to be tested in fixed genetic backgrounds (**Fig. 6E**). Thus, the sequence solutions selected *in vivo* reflected the functional preferences of the boosting immunogens, linking immunogen affinity directly to the maturation pathways that emerged following vaccination.

Together, these findings demonstrate that sequential immunization can selectively drive functional mutations favored by individual boosting immunogens and generate SF-like antibodies with broad heterologous neutralization. Importantly, the most broadly active elicited antibody followed a distinct maturation pathway from SF10, demonstrating that vaccination can access alternative molecular solutions to broad SF recognition rather than requiring precise recapitulation of a naturally evolved bnAb lineage.

## DISCUSSION

Sequential immunization and germline-targeting vaccine strategies have established that defined bnAb precursors can be activated and driven to breadth by vaccination in humans and non-human primates^18–21^. However, efficiently co-evolving complementary bnAb lineages within a polyclonal response remains an unsolved challenge and identification of bnAbs with more parsimonious pathways to vaccine elicitation would aid in development of a multi-epitope vaccine^18–21^. Here, we establish the HIV-1 silent face (SF) as an additional target for bnAb-directed vaccination and illustrate how defining functional requirements for antibody breadth can inform rational selection of more efficient maturation pathways. By integrating structural analysis, antibody minimization, and *in vivo* vaccination studies, we show that the extensive maturation of SF10 can be reduced to a limited set of heavy chain mutations that can be targeted through a rapid sequential immunization regimen. These findings extend our previous identification of the SF as a target of naturally occurring human bnAbs by demonstrating that key functional features underlying SF-directed breadth can be induced and guided through vaccination.

Previous studies of HIV-1 bnAbs, particularly those targeting the CD4 binding site, have shown that the somatic hypermutation present in mature antibodies can substantially exceed that required for broad neutralization, motivating the development of minimally mutated bnAb variants as potentially more attainable vaccine endpoints^22–24^. Our findings extend this concept to the SF, where the maturation requirements underlying breadth have remained poorly defined. Despite extensive somatic mutation present in mature SF10, a restricted set of heavy chain mutations preserved most of its breadth and potency, including several improbable mutations that could pose barriers to elicitation. Structural comparison with the clonally related SF12 antibody further revealed convergence on a shared architecture for SF recognition despite substantial sequence divergence and distinct local solutions for glycan accommodation. Thus, broad SF recognition appears to depend on conservation of key structural features without strict dependence on a single antibody sequence. This distinction is important for vaccine design because it suggests that productive SF maturation may be accessible through multiple sequence trajectories that achieve key functional features rather than requiring re-creation of the extensive mutation present in a naturally evolved bnAb.

We leveraged these defined requirements to select boosting immunogens capable of imposing specific pressures on SF10VHrev B cells. Previous mutation-guided approaches, including studies of the V3-glycan bnAb DH270.6, have proposed that improbable mutations can represent rate-limiting steps in bnAb development and that immunogens with appropriate affinity gradients can favor B cells that acquire these substitutions^4,25–28^. Whereas our engineered B41 GT1.2 prime enabled efficient engagement of SF10VHrev precursors, native Env trimers bound the precursor relatively weakly but showed increased affinity for variants containing specific breadth-associated mutations. We therefore used native Envs as selective affinity gates during boosting, such that B cells acquiring key improbable mutations that improved recognition would gain a selective advantage. Consistent with this strategy, concurrent selection of multiple improbable mutations was possible using rationally selected native-like Env immunogens, in contrast to engineered immunogens used previously^26,28^. Particularly notable was selection of the improbable S54D mutation despite the absence of a canonical AID hotspot, as well as enrichment of S31L following boosting with Envs that preferentially recognized this substitution. Moreover, different Env strains favored distinct mutational outcomes despite similar baseline affinity for SF10VHrev. These findings suggest that naturally occurring Env diversity can impose distinct selective pressures on antibody evolution that can be exploited to favor functional features associated with breadth. Such affinity-gated selection may provide an alternative to engineering every intermediate immunogen *de novo* and could help simplify maturation strategies for bnAb elicitation.

The functional consequences of this selection were apparent after a short immunization regimen. Heterologous serum neutralization emerged after only the prime and first native Env boost, and all selected monoclonal antibodies tested neutralized heterologous viruses on a multiclade panel, directly linking the observed maturation of SF10VHrev B cells to antiviral activity. Importantly, vaccination did not simply reproduce the SF10 pathway. This was exemplified by 14.1, which neutralized more viruses than mature SF10 on the tested panel despite substantially fewer VH mutations and combined features found in both SF10 and SF12. More broadly, the recurrent selection of S31L and other hydrophobic substitutions rather than SF10 S31Q demonstrated that distinct amino acid solutions can support the same desired antibody function. This sequence flexibility is consistent with the structural convergence of SF10 and SF12 and suggests that successful vaccination may not require precise reproduction of a donor-derived lineage. The existence of more tractable pathways to antibody function has been hypothesized for other bnAbs^29^, but here we demonstrate that these pathways can be reproducibly accessed and selected via vaccination.

These findings extend the concept of bnAb minimization by linking simplified antibody endpoints to immunogen driven selection *in vivo*. Natural infection provides one successful evolutionary trajectory to a mature antibody, while minimization can distinguish features necessary for function from mutations accumulated during its broader history. Immunogens can then be selected to favor B cells approaching these desired states without prescribing the exact sequence used to achieve them. This flexibility may be particularly valuable as HIV-1 vaccine development moves toward coordinating multiple bnAb specificities within polyclonal responses, where highly prescribed and lengthy immunization regimens for each individual lineage could become impractical. SF-directed bnAbs are attractive candidates for multi-epitope HIV-1 vaccine strategies because they can achieve substantial breadth and potency while providing neutralization coverage complementary to established bnAb classes^15^. Importantly, studies of polyclonal responses in an elite neutralizer have further shown that even SF-directed antibodies with more limited individual breadth can contribute alongside antibodies targeting other Env epitopes to achieve broad and potent serum neutralization^13^. Thus, vaccine induction of SF bnAbs offers the potential to generate potent and broadly neutralizing responses against a distinct Env epitope, while SF antibodies spanning a range of neutralization breadth may also enhance the coverage of a polyclonal response. Future studies will need to determine whether SF precursors can be activated and matured alongside B cells targeting complementary Env epitopes and how immunogen modifications or selective pressures introduced at one epitope influence responses directed toward another.

However, several limitations remain. SF10VHrev was designed to isolate VH maturation and therefore retains the mature SF10 CDRH3 and light chain, while the IgKI model provides a non-physiologic precursor frequency without competition from a fully polyclonal B cell repertoire. Our structural and functional analyses provide a rationale for this design, as the major experimentally defined maturation barriers were concentrated within the heavy chain, whereas the light chain exhibited greater structural plasticity and required only a limited number of probable mutations. Nevertheless, the present study does not establish that fully germline SF precursors can be efficiently activated and matured by vaccination. Future efforts to develop immunogens capable of engaging fully germline SF precursors and evaluating them in an adoptive transfer model that matches the physiologic precursor frequency and enables competition with the naïve repertoire will be important. Native Env strains recognized by minimally mutated SF10 variants provide potential starting points for these efforts and raise the possibility that naturally occurring native-like Env strains could be further exploited to simplify future immunization regimens^20,30^. Larger neutralization panels, structural characterization of elicited antibodies, and extension to SF bnAbs derived from additional germlines will also be necessary to establish the breadth and generality of the maturation pathways identified here.

Together, our findings move the silent face from a site of vulnerability defined by infection-derived bnAbs toward an experimentally accessible vaccine target. Structure- and function-guided immunogen selection enabled activation of an SF precursor, enrichment of mutations associated with breadth, and rapid induction of heterologous neutralizing responses. The emergence of broad antibodies through pathways distinct from mature SF10 further suggests that vaccination can exploit a landscape of productive maturation solutions rather than reproduce a single naturally evolved lineage. Expanding this principle to the SF and other complementary Env vulnerabilities may increase the repertoire of bnAb specificities that can ultimately be combined to generate broader protective antibody responses against HIV-1.

## MATERIALS AND METHODS

### Protein cloning, expression, and purification

All recombinant gp120 core proteins were cloned into a mammalian expression vector with an N-terminal signal peptide and a C-terminal AviTag followed by a polyhistidine tag. Antibodies were expressed in a human IgG1 backbone via co-transfection of heavy and light chain DNA at a 1:1 ratio. DS-SOSIP Envelope constructs were expressed as previously described in Joyce *et al* and co-expressed with furin at a 4:1 SOSIP:furin ratio^31^. Constructs were transfected in Expi293F cells and expressed for 4 days at 37°C. For biotinylated constructs, cells were co-transfected with a plasmid encoding for the *Escherichia coli* biotin ligase (BirA) obtained from the laboratory of K. Zinn (California Institute of Technology). Recombinant proteins were isolated from cellular supernatants and purified by affinity chromatography using either a HisTrap HP column (Cytiva, gp120 cores) HiTrap MabSelect column (Cytiva, antibodies), or PGT145 immunoaffinity column, followed by size exclusion chromatography on Superdex 200 Increase 10/300 GL (Cytiva) or Superose 6 200pg (Cytiva) columns using an AKTA pure 25 M1 system (Cytiva). All proteins were stored in 1× TBS-Az at 4°C. Detailed sequence and design information for all recombinant viral protein constructs used in this study can be found in Table S1.

### Cryo-EM sample preparation

Quantifoil Cu 1.2/1.3 300 mesh grids were glow-discharged for 60 s at 10 mA (PELCO easiGlow) before sample application. For 01428 and TRO-11 structures, SOSIP (3 mg/ml) was first incubated with SF10 antibody Fab fragments at a 1:2 gp120–to–Fab molar ratio for 1 hr in 1× TBS before the addition of fiducial Fab (either PGT145 or BG18) at 1:2 molar ratio for an additional 1hr. For the SF10VHrev structure, SF10VHrev was added at a 1:2 gp120-to-Fab molar ratio and incubated overnight at 4°C before the addition of PGT145 for 2hr at 4°C. Samples were separated from free Fab using a Superose 6 200pg (Cytiva) column. Grids were blotted for 3 s at room temperature, 100% humidity, and blot force 0; plunge-frozen in liquid ethane; and stored in liquid nitrogen until data collection.

### Cryo-EM data collection and processing

Single-particle cryo-EM datasets were collected on either a Titan Krios transmission electron microscope (Thermo Fisher Scientific) equipped with a K3 direct electron detector and energy filter (Gatan), operating at 300 kV for the 01428-SF10-BG18 structure or a Glacios transmission electron microscope (Thermo Fisher Scientific) equipped with a K3 direct electron detector operating at 200kV for the TRO-11 and B41 GT1.2 structures. For the 01428 structure, movies were acquired in an automated fashion using SerialEM v3.8 and a 3-by-3 multishot data acquisition pattern, with a total dose of 60 electrons/Å^2^ accumulated across 40 frames with a calibrated pixel size of 0.86 Å and a defocus range of −1 to −2.5 μm. For the TRO-11-SF10-PGT145 and B41 GT1.2-SF10VHrev-PGT145 structures, movies were acquired in an automated fashion using Thermo Fisher EPU with a total dose of 60 or 40 electrons/Å^2^ respectively accumulated across 40 frames and with a calibrated pixel size of 1.2 Å and a defocus range of -1 to -2.5 µm. For all datasets, motion correction, contrast transfer function (CTF) estimation, micrograph curation, reference-free particle picking, and particle extraction as well as iterative 2D classification and ab initio modeling followed by heterogenous reconstruction were carried out using cryoSPARC v4.1^32^. Particles corresponding to SOSIP–Fab complexes subsequently underwent iterative 3D classification in Relion (k = 4), followed by re-extraction without binning and nonuniform refinement with C1 symmetry in cryoSPARC^33,34^. For the 01428 structure, further structure refinement was performed in cryoDRGN to remove trimers with sub-stoichiometric Fab binding^35^. Resolution was estimated according to the 0.143 Fourier shell correlation (FSC) cutoff criterion in cryoSPARC. Further details of data collection and processing can be found in Table S2.

### Cryo-EM structure modeling, refinement, and analyses

Initial models the 01428 and TRO-11 trimers were generated in Alphafold while a previously solved B41 structure (PDB ID: 6OKP) was used for B41 GT1.2^15,36^. Initial models were docked into sharpened global nonuniform maps using ChimeraX, with unsharpened maps used for glycan building^37^. Structures were then manually corrected in Coot and refined using iterative rounds of refinement in Phenix and manual building in Coot^38,39^. Glycans were modeled at potential *N*-linked glycosylation sites using the carbohydrate module in Coot. Validation of refined model coordinates was performed in MolProbity^40^. UCSF ChimeraX v1.8 was used for structure visualization^37^. Buried surface areas were calculated using ChimeraX1.8 and a 1.4-Å probe. Potential hydrogen bond (less than 4 Å and with an A-D-H angle greater than 90°) and van der Waals interaction assignments were performed using ChimeraX. Antibody residues were denoted by Kabat numbering^41^.

### X-ray crystallography data collection, refinement, and analysis

Crystallization was carried out at room temperature using the sitting drop vapor diffusion method by mixing equal volumes of Fab and reservoir using a TTP LabTech Mosquito robot and commercially-available screens (Hampton Research). Crystals were obtained in 25%w/v polyethylene glycol, 0.1M TRIS and 0.2M Lithium sulfate (SF10 Fab: 38RE) and 1.76M Sodium di0hydrogen phosphate, 0.036M di-Potassium hydrogen phosphate (SF10VHrev: 38RF). Crystals were cryoprotected in a mixture of well solution and 25% glycerol and cryopreserved in liquid nitrogen. X-ray diffraction data were collected at the Stanford Synchrotron Radiation Lightsource (SSRL) beamline 14-1 on a Pilatus 6M pixel detector at a wavelength of 1.19Å (38RE) and beamline 12-2 on a Dectris Eiger X 16M pixel detector at a wavelength of 0.979 Å (38RF). Data from single crystals of 38RE and 38RF were indexed and integrated in XDS and merged using AIMLESS v0.7.4 in *CCP4* v7.0.6. The structures for both 38RE and 38RF were determined by molecular replacement in Phenix v1.21.2 using the coordinates for SF12 HC (PDB 6OKQ) and 1269 LC (PDB 6B0A) and refined using Phenix and cycles of manual building in Coot v0.9.6. Details of crystallographic refinement can be found in Table S3.

### Enzyme-linked immunosorbent assays

ELISAs of SF10 light chain reversions and B41 GT1.2 with SF10VHrev in Figure S3 were performed similarly to Rubio et al^42^. Briefly, ELISAs were performed by immobilizing viral antigens on streptavidin-coated 96-well assay plates (Corning) at a concentration of 2 μg ml^−1^ in 1× TBS-Az for 1 hour at 22°C. After incubation, plates were washed 3 times with 200µL washing buffer (1× TBS plus 0.05% Tween-20; TBS-T). After washing the plates with TBS-T washing buffer, serial dilutions of mAbs were added to the plates at a top concentration of 100 μg ml^−1^ in TBS-TMS (TBS plus 0.05% Tween-20, 1% nonfat dry milk, and 1% goat serum) blocking buffer and incubated for 30 min at room temperature. Plates were washed in TBS-T and incubated with horseradish peroxidase (HRP)–conjugated goat anti–human Ig Fc antibody (Southern Biotech, 2047-050) at a dilution of 1:4000 in TBS-TMS for 30 min at room temperature. After washing the plates with TBS-T, 100 μl of 1-Step Ultra TMB-ELISA substrate (Thermo Scientific, 34029) was added to each well for 3 min. Reactions were quenched with 100 μl per well of a 1 M HCl solution, and absorbance at 450 nm was measured with a Tecan Infinite M Plex microplate reader using iControl 2.0 software. Data obtained were plotted with GraphPad Prism 10. For statistical comparisons, ELISAs were completed in biological triplicate with two technical replicates on each plate. Area under the curve was fit for each experiment and statistics were calculated over all replicates.

ELISAs with gp120c B41GT1.2, B41GT1.2 KO, B41KO, 001428 KO and CNE8 KO and native Env gp120c B41, 001428, CNE8, CNE55 and Tro11 were performed as previously described^43^. Briefly, 96-well plates were coated with 50μl of a solution of the trimer at 2μg ml^−1^ in 1× PBS (Corning, 20-030-CV) and incubated at 4°C overnight. Plates were washed with washing buffer (1× PBS with 0.05% Tween-20; Sigma-Aldrich, P7949) and blocked (1× PBS with 4% milk) for 1h at room temperature. Serum samples were assayed at a 1:60 starting dilution and the antibody controls were added at 5μg ml^−1^ all followed by seven additional threefold serial dilutions for 1h at room temperature. Plates were washed three times and incubated with an anti-human IgG secondary antibody conjugated to horseradish peroxidase (HRP; Jackson ImmunoResearch, 115-035-071) or anti-mouse secondary antibody (Peroxidase AffiniPure® Goat Anti-Mouse IgG, Fcγ fragment specific 115-035-071) at a 1:5,000 dilution in washing buffer for 1h at room temperature. Plates were developed by adding ABTS, and optical density was measured at 405nm. AUC data were calculated and plotted with GraphPad Prism 10.

### IgG purification from mouse serum

Serum samples were heat-inactivated for 1 hour at 56 °C. Polyclonal IgGs were purified from serum by incubation with an equal volume of Protein G Sepharose resin (GE Healthcare) at 4 °C with overnight shaking. The resin was washed with PBS and IgGs were eluted using 0.1 M glycine (pH 2.5). Eluates were buffer exchanged into PBS using 30 kDa Amicon Ultra centrifugal filter units (MilliporeSigma).

### Biolayer interferometry

BLI measurements were performed on an Octet Red96 system (FortéBio) at 25°C in Octet buffer (TBS plus 0.1% bovine serum albumin and 0.05% Tween-20). For assays used to calculate binding kinetic data, IgGs were immobilized onto protein A (ProA) biosensors (Sartorius) and dipped into a concentration series of monomeric analytes (top concentrations ranging from 1 to 10 μM) for 90s and allowed to dissociate in Octet buffer for 180s. For assays using ProA biosensors (Sartorius), regeneration of biosensors was achieved using 10 mM glycine, pH 1.5, and neutralized with Octet buffer. Kinetic analyses were used after the subtraction of reference curves to derive kon and koff rates and binding constants (*K*D) using a 1:1 binding model in the FortéBio Data Analysis 9.0 software and GraphPad Prism 10 for monomeric affinities. For kinetic analysis of the single cycle kinetics experiment, B41 MD39 WT or B41 MD39 GT1.2 SOSIP was immobilized on a ProA tip loaded with PGT145 IgG before being dipped into successive concentrations of SF10VHrev Fab for 120s with 60s dissociation steps between each association before a final dissocation of 600s. This curve was fit using the single cycle kinetics function in the ANABEL R package using RStudio^44^. BLI experiments with multivalent nanoparticle immunogens from Supplemental Figure 3 were performed by loading SF10VHrev IgG onto a ProA biosensor before association with 1µM of nanoparticle immunogen.

### Yeast display

Yeast libraries were generated in the pCTCON2 backbone. Briefly, for the minSF10 scFv library, a linear DNA insert bearing the minSF10 sequence in scFv format (Nterm-LC-GSlink-HC-Cterm) with a 3x GGGGS linker was ordered from Integrated DNA Technologies. The insert was subsequently combinatorially mutagenized back to the mature amino acid codon at each site reverted in minSF10 (29 sites with two possible amino acids for a 5.34x10^8^ theoretical library size). This was accomplished using overlap extension PCR with a mix of forward and reverse mutagenesis primers, with three total rounds of mutagenesis performed. The mutagenized insert was stitched together using a final PCR reaction before being gel extracted, sequenced to confirm mutated sites, and amplified using PCR to produce at least 10µg linear insert for library transformation. The linearized pCTCON backbone was generated using standard PCR methods.

The B41 GT1.2 library was designed using the SF10-01428 structure. Surface exposed sites located near the CDHR1 of SF10 were selected for targeted mutagenesis (positions 293, 295, 444, and 446). NNK codons were used to produce random changes at the desired sites while reducing introduction of premature stop codons (theoretical library size 160,000). Mutagenesis and generation of linear insert for transformation was accomplished using overlap extension PCR in a similar manner to the scFv libraries.

Library transformation and magnetic activated cell sorting was done as described in Gristick et al^45^. The notable difference was the use 25µg of linear insert DNA and 5µg of backbone for the B41 library. Serial dilutions of libraries were plated after transformation to confirm adequate transformation efficiency to represent library diversity. To prepare the scFv yeast libraries for FACS analysis, cells were pelleted at 3000 rpm for 2 min and washed twice with PBSF. Cells in excess of 10x of library diversity were stained at a density of 10^7^ cells/ml with 1:1000 anti-c-Myc antibody conjugated to FITC (Abcam, ab1394) and 1 µM biotinylated gp120c (either B41, 01428, or CNE55) precomplexed with Streptavidin-AF-647 (ThermoScientific, S32357) and incubated for 1h on a rotator at 4°C. Cells were then washed twice with PBSF and analyzed on a MACSQuant Analyzer (Miltenyi Biotec) or sorted using an SH800 cell sorter system (Sony). Cells that stained double-positive for both c-Myc and gp120c-Strep-AF-647 were collected and grown in 5 mL SDCAA-carb for 1-2 days at 30°C and 250 rpm and then transferred to 100 mL SDCAA-carb for an additional 1-2 days at 30°C and 250 rpm. After each round of sorting, DNA was extracted for pooled Illumina sequencing. Gp120c libraries from each clade were independently sorted between 3-5 times to enrichment prior to Illumina sequencing to identify enriched mutations. Mutations that were enriched in one or more libraries were used to produce minSF10.20.

To prepare the gp120c yeast libraries for FACS analysis, cells were pelleted at 3000 rpm for 2 min and washed twice with PBSF. Cells were then stained at a density of 10^7^ cells/ml with 1:1000 anti-c-Myc antibody conjugated to FITC (Abcam, ab1394) and 1 µM SF10VHrev and incubated for 1h on a rotator at 4°C. Cells were then washed twice with PBSF and resuspended in PBSF with 1:1000 goat anti-human antibody conjugated to AlexaFluor647 (Invitrogen, A-21445) and incubated for 30 min at 4°C. Cells were then washed twice more with PBSF and analyzed on a MACSQuant Analyzer (Miltenyi Biotec) or sorted using an SH800 cell sorter system (Sony). Cells that stained double-positive for both c-Myc and SF10VHrev were collected and grown in 5 mL SDCAA-carb for 1 - 2 days at 30°C and 250 rpm and then transferred to 100 mL SDCAA-carb for an additional 1-2 days at 30°C and 250 rpm. After each round of sorting, DNA was extracted for pooled Illumina sequencing. Gp120c libraries were sorted 3 times and final candidate immunogens were selected from enriched sequences identified from the third sort.

### Nanoparticle expression and characterization

SpyCatcher003-mi3 nanoparticles were expressed and purified as previously described^46^.SpyCatcher003-ferritin was expressed and purified as described in Wang et al^47^. Briefly, both nanoparticles were expressed in Escherichia coli BL21 DE3 cells before bacterial protein purification and were separated from aggregates and non-full size nanoparticles using size exclusion chromatography with a Superose 6 200pg (Cytiva) column. Mi3 eluted in the void volume of this column and ferritin nanoparticles eluted around 12.5mL. Purified nanoparticles were characterized using negative stain electron microscopy and dynamic light scattering to confirm expected size distribution. Samples used for animal immunizations further underwent endotoxin removal using Triton X114 as described^46^. Excess Triton X114 was removed using an overnight incubation with BioBeads SM-2 (BioRad) followed by 5min centrifugation at 14,000x g to remove the beads. Conjugation was performed overnight at room temperature using 2:1 gp120c-to-subunit ratio for the priming immunogen and a 1:2 protomer-to-subunit ratio for SOSIP-nanoparticle conjugates. Nanoparticles were separated from free antigen using a Superose 6 200pg column before being stored in endotoxin-free PBS (Sigma Aldrich) at 4°C prior to immunization.

### Pseudo-virus neutralization assays

The TZM-bl neutralization assay was performed as previously described^48^. Briefly, mAbs or purified IgG were serially titrated in duplicate wells using a primary concentration of 50 mg/ml (mAbs) or 150 mg/ml (purified IgG) and a 5-fold dilution series. Following addition of HIV-1 Env pseudovirus, plates were incubated for 1 hour at 37° C, followed by addition of TZM.bl cells (1x10^4^/well) in the presence of 11ug/ml DEAE-Dextran (Sigma). Wells containing cells + pseudovirus (without sample) or cells alone acted as positive and negative infection controls, respectively. Following a 48 hour incubation, plates were harvested using Promega Bright-Glo luciferase reagent (Madison, WI) and luminescence detected with a Promega GloMax luminometer. Titers are reported as the concentration of mAb that inhibited 50% or 80% virus infection (IC50 and IC80 titers, respectively). Virus pseudotyped with the envelope protein of murine leukemia virus (MuLV) was used as negative control. Additional information on this assay and supporting protocols may be found at: http://www.hiv.lanl.gov/content/nab-reference-strains/html/home.htm

### Mice

Mice expressing the precursor of the human SF10 bnAb were produced in-house on the C57BL/6 strain background. All animal procedures were performed in accordance with protocols approved by The Wistar Institute IACUC. All mouse cages were on ventilated, ducted single-use racks. Cages were pre-bedded with Alpha-Dry bedding. Water bottles and cage supplies were irradiated. Cages were changed every 2 weeks, and lids and feeders were changed once a month inside a class-II biosafety cabinet using aseptic technique. Animals receive HCL water with a 3.0pH. The Wistar Institute’s animal facility is operated as a modified barrier and with room-level protection. Holding rooms are negative to the corridors. Up to five adult mice or two adult mice plus one female with a litter may be housed in a cage (81 square inches). Temperature is within 64 to 79 degrees Fahrenheit and humidity is within 30% to 70%. Light cycle is 12-h on starting at 6:00 and 12-h off starting at 18:00.

### SF10VHrev mouse generation and validation

Mice carrying the Ig V(D)J genes encoding the SF10VHrev antibody were produced using a CRISPR/Cas9 and Adeno Associated Virus (AAV)-based protocol (in preparation). Briefly, we used an AAV-based donor template carrying the pre-rearranged immunoglobulin genes of the partially reverted SF10 heavy-chain sequence (VHrev) together with the mature pre-rearranged SF10 light-chain genes to insert these genes into the heavy chain and light chain loci respectively. Fertilized mouse zygotes were incubated with the AAV donor template and electroporated to deliver CRISPR– Cas9 ribonucleoprotein complexes. Next day, two-cell stage embryos were implanted into pseudopregnant females. Resulting pups were screened for correct insertion of V(D)J genes by PCR using genomic DNA and were further characterized to confirm normal B cell development. Briefly, peripheral blood, spleens, and bone marrow were collected from SF10VHrev Ig KI mice. Blood samples were treated with ACK lysis buffer to remove red blood cells prior to staining. Spleens were mechanically dissociated through 70-µm cell strainers, followed by red blood cell lysis and filtration to obtain single-cell suspensions. Bone marrow was isolated from the femurs by centrifugation, followed by red blood cell lysis and filtration through 70-µm strainers. Cell suspensions were maintained on ice and stained with the antibody combinations in Table S4 respectively. B-cells were analyzed by flow cytometry.

### Mouse immunization

Male and female SF10VHrev mice of 6–8 weeks of age were equally distributed in groups for immunization experiments. SF10VHrev mice were immunized every 2 weeks intraperitoneally (i.p.) with 10μg of protein and 3 µg of SMNP as adjuvant. Serum samples were collected 2 weeks after each immunization. Mice were maintained on regular chow.

### Single B-cell sorting

Draining parathymic lymph nodes were harvested from immunized mice 2 weeks after each immunization (w2, w4 and w6, respectively) and processed into single-cell suspensions by mechanical dissociation through a 70-µm cell strainer. Red blood cells were lysed using ACK lysis buffer. Cells were washed and resuspended in FACS buffer. To select antigen-specific germinal center (GC) B cells, single-cell suspensions were stained with a viability dye and fluorophore-conjugated antibodies against surface markers, including B220, GL7, and CD95 (Fas). Antigen specificity was determined using biotinylated TRO-11gp120c and/or CNE8 gp120c probes conjugated with streptavidin-PE or -AF647 and a biotinylated silent face epitope knockout probe, TRO-11 KO gp120c, conjugated with streptavidin BV605. Silent face-specific B cells were selected as Dump^-^(Dead^-^CD4^−^CD8^−^Gr1^−^F4/80^−^) B220^+^; GL7^+^, CD95^+^; KO bait^−^; bait^+/+^. Antigen-specific GC B cells were sorted as single cells into 96-well PCR plates containing 5µl of TCL buffer with 2% B mercaptoethanol. Plates were immediately sealed, briefly centrifuged, snap-frozen on dry ice, and stored at −80 °C until further processing. cDNA from single cells was obtained by reverse transcription (Superscript III, Invitrogen) and used for amplification by nested PCR using sequencing primers listed in Table S5. PCR protocols (annealing (°C)/elongation (sec)/number of cycles): 1st PCR (IgG IgH): 52/55/50; 2nd PCR (IgG IgH):54/55/50; 1st PCR (Igκ): 46/55/50; 2nd PCR (Igκ): 50/50/50; 1st PCR (IgM IgH): 52/55/50; 2nd PCR (IgM IgH): 54/55/50.

### Donut plots

Amino-acid change classification was performed at user-defined positions in a multiple sequence alignment using a bespoke script. The alignment was supplied as codon-aligned nucleotide or amino-acid FASTA; nucleotide alignments were translated in-frame to amino acids. Positions were specified in ungapped anchor-sequence coordinates and mapped to alignment columns. For each position, residues from all sequences were compared to both the bnAb residue and the germline (GL) residue at the same column; gaps and non-standard residues were excluded. Each residue was assigned to a single category: (i) bnAb-like (exact match to bnAb), (ii) GL-like (exact match to germline), (iii) Similar (different from bnAb but in the same physicochemical class as bnAb: positive, negative, hydrophobic, polar, or special), (iv) Opposite (charge inversion relative to bnAb, positive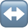negative; optionally treating charge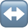hydrophobic swaps as opposite), or (v) Other change (all remaining substitutions). Counts per category were aggregated at each position and used to generate donut plots and summary tables.

## DATA AND CODE AVAILABILITY

The atomic models and cryo-EM maps generated for SF10 bound to TRO-11 and 01428 DS-SOSIPs as well as SF10VHrev bound to B41 GT1.2 DS-SOSIP, and SF10VHrev and SF10 Fabs have been deposited to the Protein Data Bank (PDB; http://www.rcsb.org/) and the Electron Microscopy Databank (EMDB; http://www.emdataresource.org/) under accession codes PDB: 38YH, 38YI and 38YJ, respectively, and EMD: -79234, -9235, and -79236, respectively. The atomic models generated via X-ray crystallography for the SF10 Fab and SF10VHrev Fab have been deposited to the Protein Data Bank (PDB) under accession codes 38RE and 38RF, respectively. Sequences for all constructs are available in Table S1. All data are publicly available as of the date of publication. This paper does not report original code. Any additional information required to reanalyze the data reported in this paper is available from the lead contact upon request.

## ACKNOWLEDGEMENTS

We thank members from the laboratory of Dr. Pamela Bjorkman (California Institute of Technology) and Dr. Peter S. Kim (Stanford) for cell lines, HIV-1 Env expression constructs, and technical assistance with pseudovirus neutralization assays and yeast display. We thank Dr. Darrell Irvine for generously providing the SMNP adjuvant used for the *in vivo* experiments in this study. Cryo-EM data for this work was collected at the Stanford-SLAC cryo-EM resource center with support from Dr. Elizabeth Montabana, Dr. Bharti Singal, and Dr. Haoqing Wang. The authors would like to thank Dr. Daniel Fernandez for his assistance with protein crystallography, which was performed at the Stanford Sarafan ChEM-H Macromolecular Structure Group Core Facility. Use of the Stanford Synchrotron Radiation Lightsource, SLAC National Accelerator Laboratory, is supported by the U.S. Department of Energy, Office of Science, Office of Basic Energy Sciences under Contract No. DE-AC02-76SF00515. The SSRL Structural Molecular Biology Program is supported by the DOE Office of Biological and Environmental Research, and by the National Institutes of Health, National Institute of General Medical Sciences (P30GM133894). The contents of this publication are solely the responsibility of the authors and do not necessarily represent the official views of NIGMS or NIH. C.O.B. is a HHMI Freeman Hrabowski Scholar. This article is subject to HHMI’s Open Access to Publications Policy. HHMI lab heads have previously granted a non-exclusive CC BY 4.0 license to the public and a sublicensable license to HHMI in their research articles. Pursuant to those licenses, the author-accepted manuscript of this article can be made freely available under a CC BY 4.0 license immediately upon publication. Finally, we thank all members of the Escolano and Barnes labs for their careful review and feedback on the manuscript.

## AUTHOR CONTRIBUTIONS

J.J.C, M.B.P, A.E, and C.O.B conceptualized and designed the project. J.J.C, J.J.N, and C.O.B collected and analyzed cryo-electron microscopy data. O.O. and C.O.B. collected and analyzed x-ray crystallography data. J.J.C and M.B. performed the yeast display selections. J.J.C. generated and tested immunogens and nanoparticles. Y.E.L, I.R, M.R.E, J. P., and T.C. aided in expression and purification of protein constructs used in this study. M.B.P, S.M. and M.G.G. generated and characterized the immunoglobulin knock in mice, maintained the mouse colony, carried out animal immunizations, and performed B cell sorting experiments. J.J.C and M.B.P. analyzed elicited mouse antibody sequences and J.J.C. expressed and characterized elicited monoclonal antibodies. M.S.S. carried out pseudovirus neutralization assays. J.J.C, M.B.P, A.E, and C.O.B visualized the data, wrote the original draft and received review and editing from all authors. A.E. and C.O.B. obtained funding and supervised the project.

## SUPPLEMENTARY MATERIALS

**Supplementary Figure 1.**
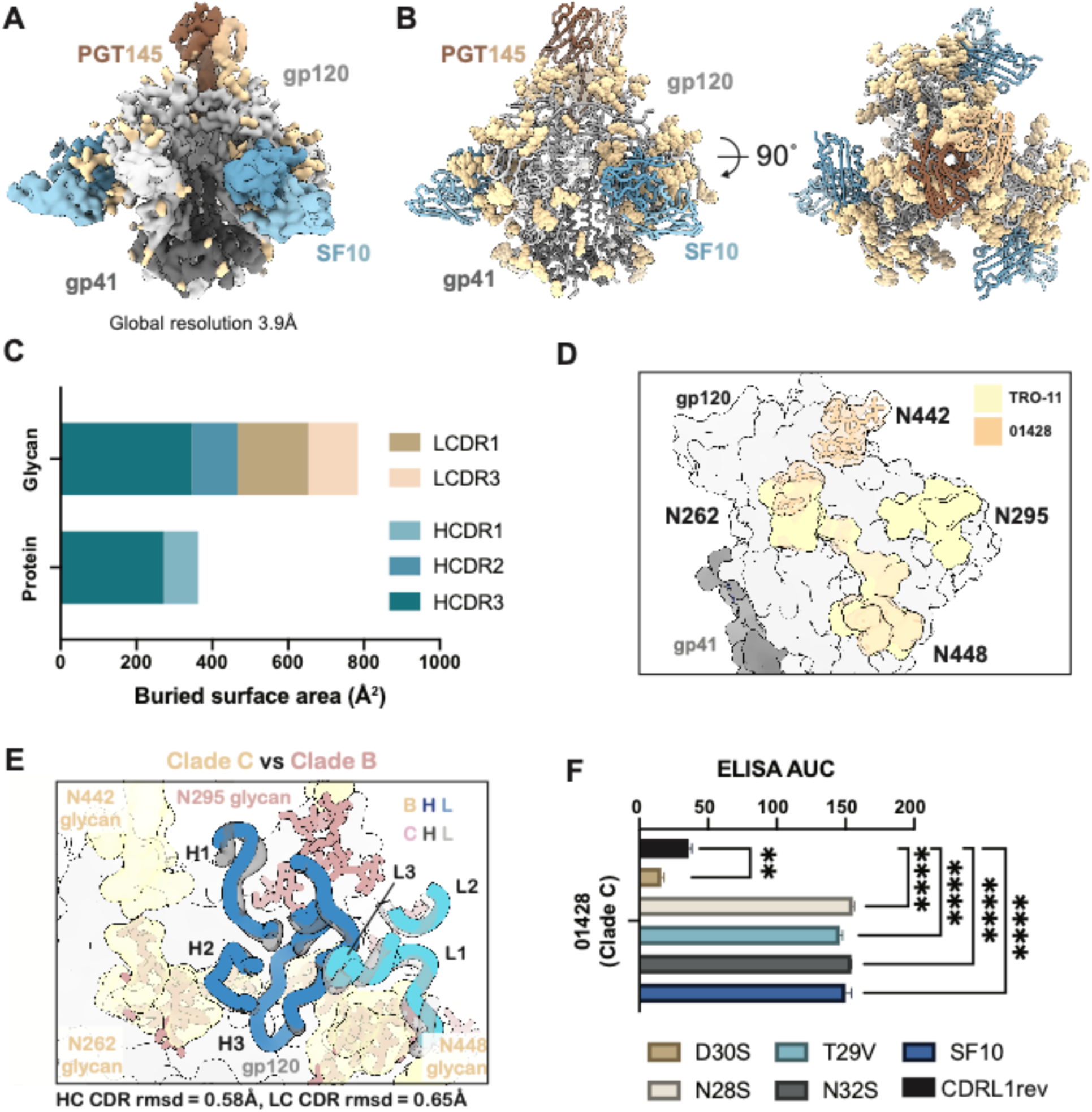
Comparison of SF antibody cryo-EM structures reveals conserved angle of approach despite altered glycan landscape and positioning. A,B) Structural density and model of SF10-PGT145-TRO-11 structure. **C)** Epitope buried surface area for SF10 bound to 01428. **D)** Comparison of the N262, N295, N442, and N448 glycans in clade B TRO-11 and clade C 01428 structures. Clade C glycans are shown as sticks and transparent for comparison. **E)** Superposition of SF10 CDR loops from TRO-11 (blue) and 01428 (black) structures. Glycans are colored as depicted in the legend and root mean square deviation of CDR loops is shown below. **F)** ELISA AUC of SF10 CDRL1 reversion variants binding to 01428 gp120c. Statistical analysis was performed using one-way ANOVA comparing AUC to to a CDRL1rev control condition with Bonferroni correction for multiple hypothesis testing.

**Supplementary Figure 2.**
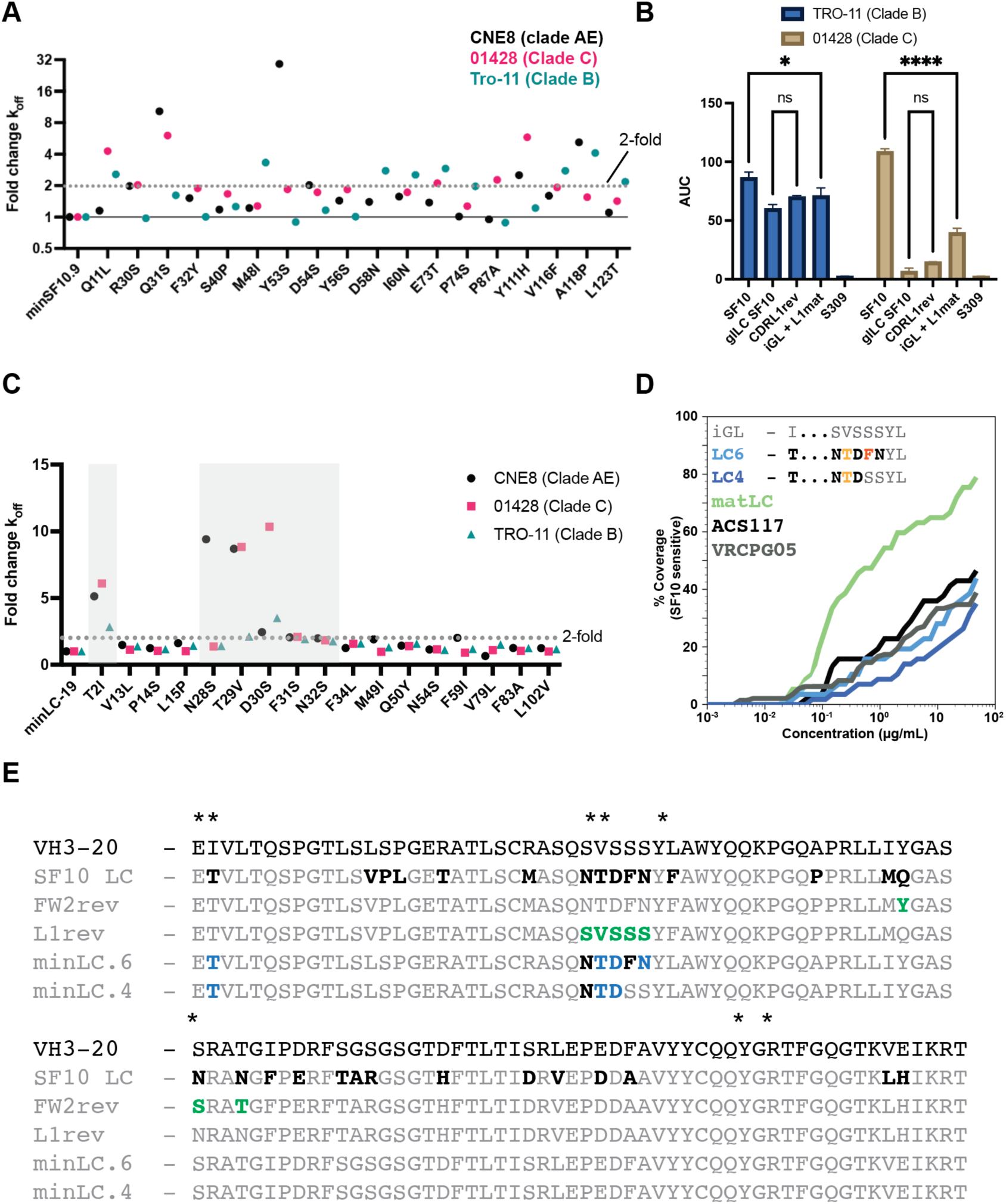
Minimization of the SF10 light chain highlights a critical role for the CDRL1-mediated N448 glycan contact site. **A)** Supporting data for minimization of the SF10 heavy chain. Fold change *k_off_* is calculated for the reversion of the mutation shown relative to the minSF10.20 baseline. **B)** ELISA data for reverted light chain variants binding to gp120c. Significance was determined from n=3 replicates using 2-way ANOVA and Sidak’s multiple comparisons test. ns p≥0.05, ** p<0.01, **** p<0.0001. **C)** Fold change *k_off_* highlighting key role of somatic mutations contacting the N448 glycan in the light chain. **D)** Neutralization data summarizing two minimized light chain variants (LC.6 and LC.4) containing intact N448 glycan contact sites paired with minSF10.11 heavy chain. Variants were tested on the 65 SF10-sensitive strain panel. Neutralization of ACS117 and VRC-PG20 are shown for comparison and sequences of the relevant contact residues are highlighted in the panel. Despite containing only 4 light chain mutations, minLC.4 retains neutralization breadth across multiple clades of HIV-1. **E)** Sequence alignments of light chain variants tested. Residues contacting the N448 glycan are denoted with an asterisk. Mutations are highlighted in bold.

**Supplementary Figure 3.**
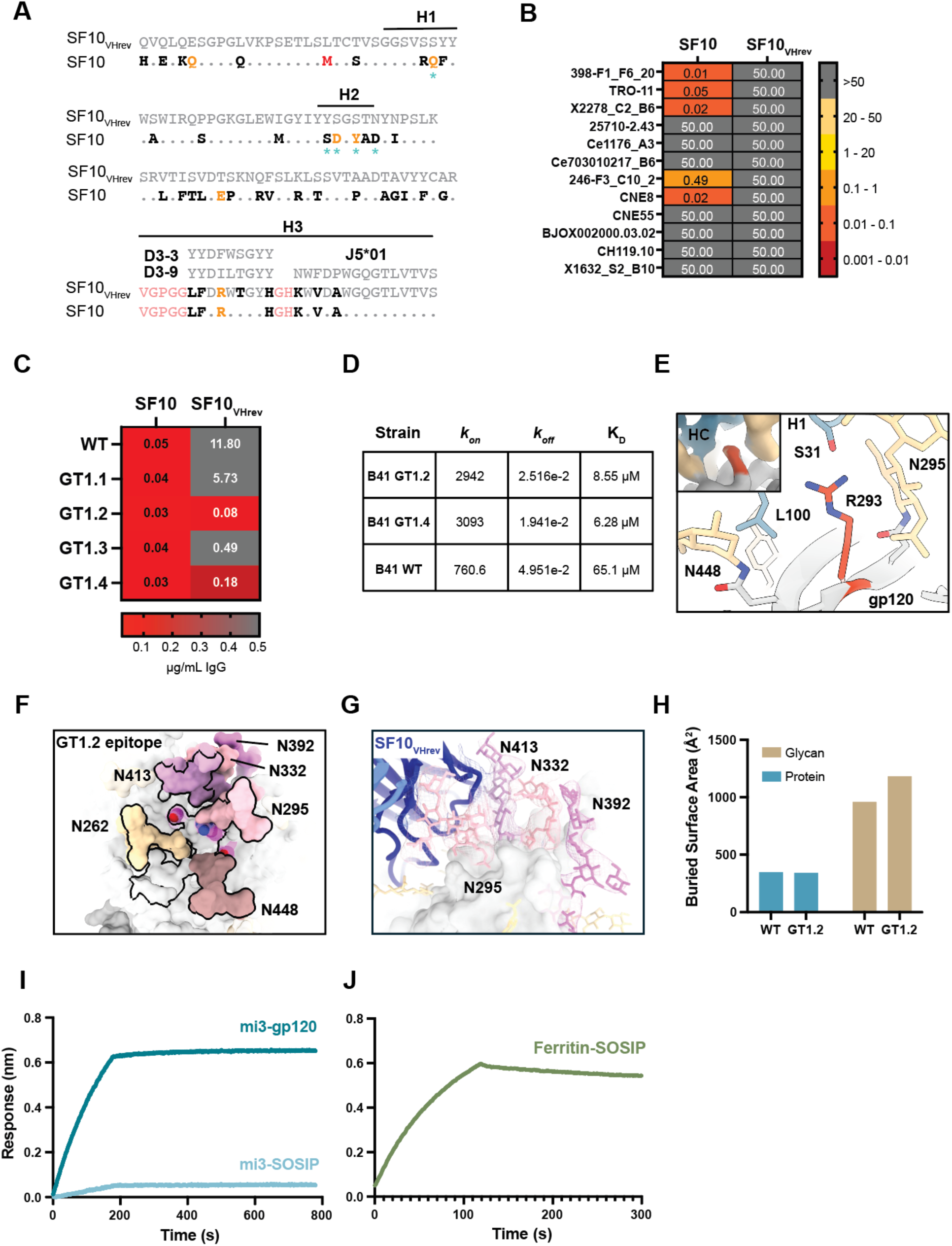
Engineering and characterization of a SF10_VHrev_-targeted protein immunogen. **A)** Sequence alignment of SF10 and SF10_VHrev_ with reverted mutations highlighted. **B)** Neutralization of the 12-strain global panel by SF10 and SF10_VHrev_. **C)** ELISA EC50 for SF10_VHrev_ binding to candidate gp120c immunogens selected via yeast display. **D)** Kinetic parameters for B41, B41 GT1.2, and B41 GT1.4 gp120c immunogens determined by BLI. Parameters were fit globally over 4 independent concentrations. **E)** Structural environment of the shared E293R mutation, with supporting density from the B41 GT1.2 cryo-EM structure in a pullout. **F)** Epitope of B41 GT1.2 is highlighted in black line, with glycans colored varying shades of pink. **G)** Glycan network observed in B41 GT1.2 with supporting density shown as mesh. This network results in a 23% increase in buried surface area, as demonstrated in panel **H). I)** BLI curves for SF10_VHrev_ IgG binding to 1µM of mi3-B41 GT1.2 gp120c or 1µM of mi3-B41 GT1.2 SOSIP. **J)** BLI curve for SF10_VHrev_ IgG binding to 1µM ferritin-SOSIP conjugated at 1:2 molar ratio (SOSIP:ferritin).

**Supplementary Figure 4.**
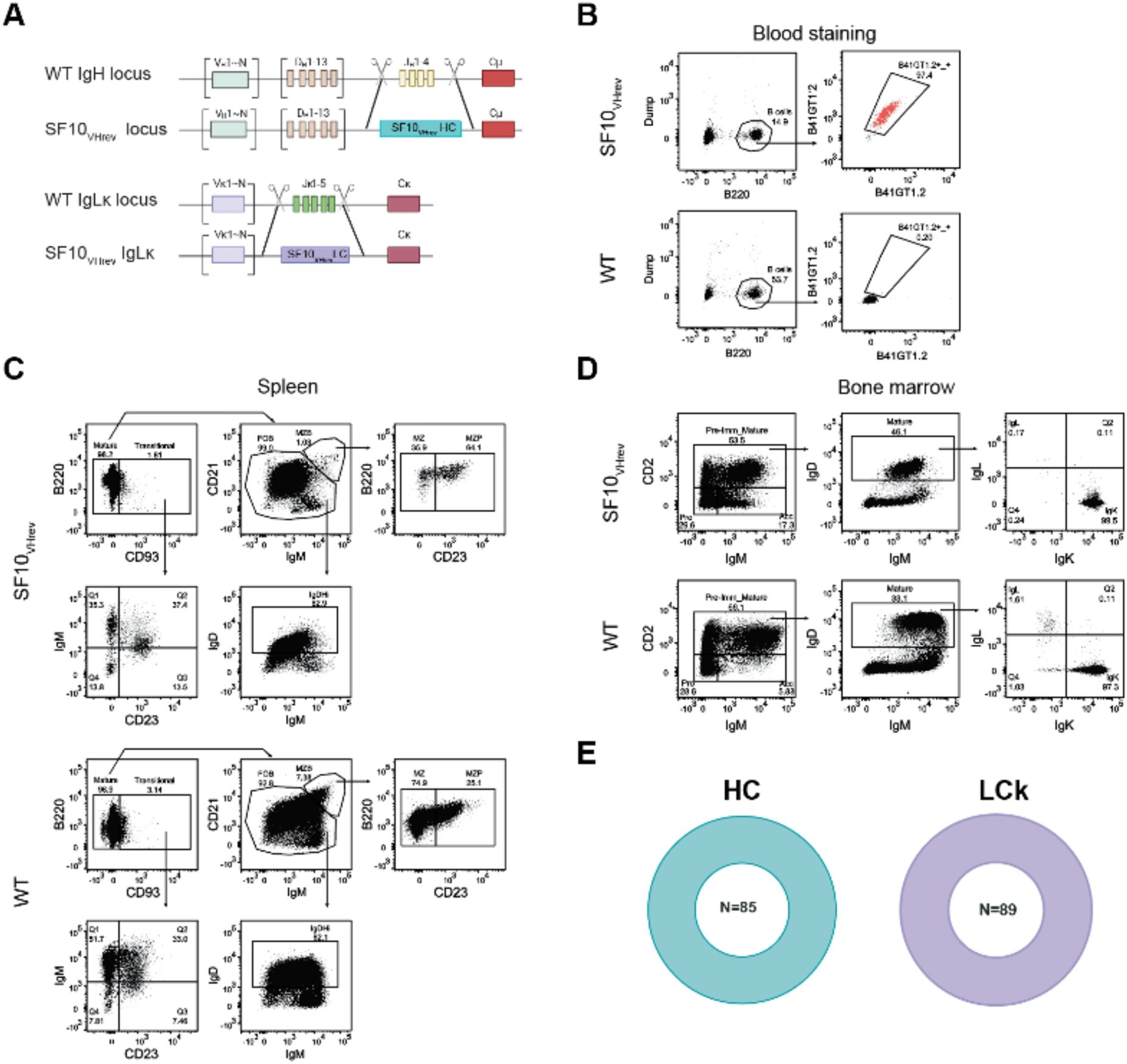
Characterization of SF10V_rev_ Ig KI mice. **A)** Diagram illustrating the transgenesis strategy to genetically engineer the mouse IgH and IgK loci. **B**). Representative flow cytometric analysis of B41GT1.2-specific B-cells in the blood of wild-type (WT) and SF10VH_rev_ Ig KI mice. **C,D**). Representative analysis of B-cell populations in the spleen (**C**) and bone marrow (BM) (**D**) of WT and SF10VH_rev_ Ig KI mice. Viable, single-cell, size-gated B-cells were separated from non-B cells by gating on B220^+^ Dump^−^ (**C**) Pre-gated splenic B-cells were further gated as Mature B-cells (B220^+^, CD93^-^) and transitional B-cells (B220^+^, CD93^+^). Mature B-cells were further gated into Follicular B cells (FOB) (CD21^lo^, IgM^+^), and marginal Zone B-cells (MZB) (CD93^−^, CD21^hi^, IgM^hi^). From them, MZB cells were further gated into marginal zone B-cell precursors (MZP) (CD23^+^) and mature MZ (CD23^-^). FOB cells were further gated into follicular I and II (Fol I/II) (CD23^−^, IgD^hi^). Transitional B-cells (B220^+^, CD93^+^) were further gated into T1(IgM^hi^, CD23^-^), T2 (IgM^hi^, CD23^+^), T3 (IgM^lo^, CD23^+^). (**D**), Pre-gated BM B-cells were classified as Pro B-cells (CD2^-^, IgM^-^), Pre-immature/mature B-cells (CD2^+^, IgM^lo/hi^) and Accelerated B-cells (Acc) (IgM^+^, CD2^-^). Pre-immature/mature B-cells were further gated into Mature (IgD^hi^, IgM^+^) and from these, B-cells were gated according to their use of lambda (IgL^+^) or kappa light chains (IgK^+^). **E**) Pie charts show the frequency of single B-cells expressing the SF10VH_rev_ KI VDJ and VJ genes in naïve SF10VH_rev_ Ig KI mice. The number in the middle of the pie chart corresponds to the number of naïve B-cells analyzed.

**Supplementary Figure 5.**
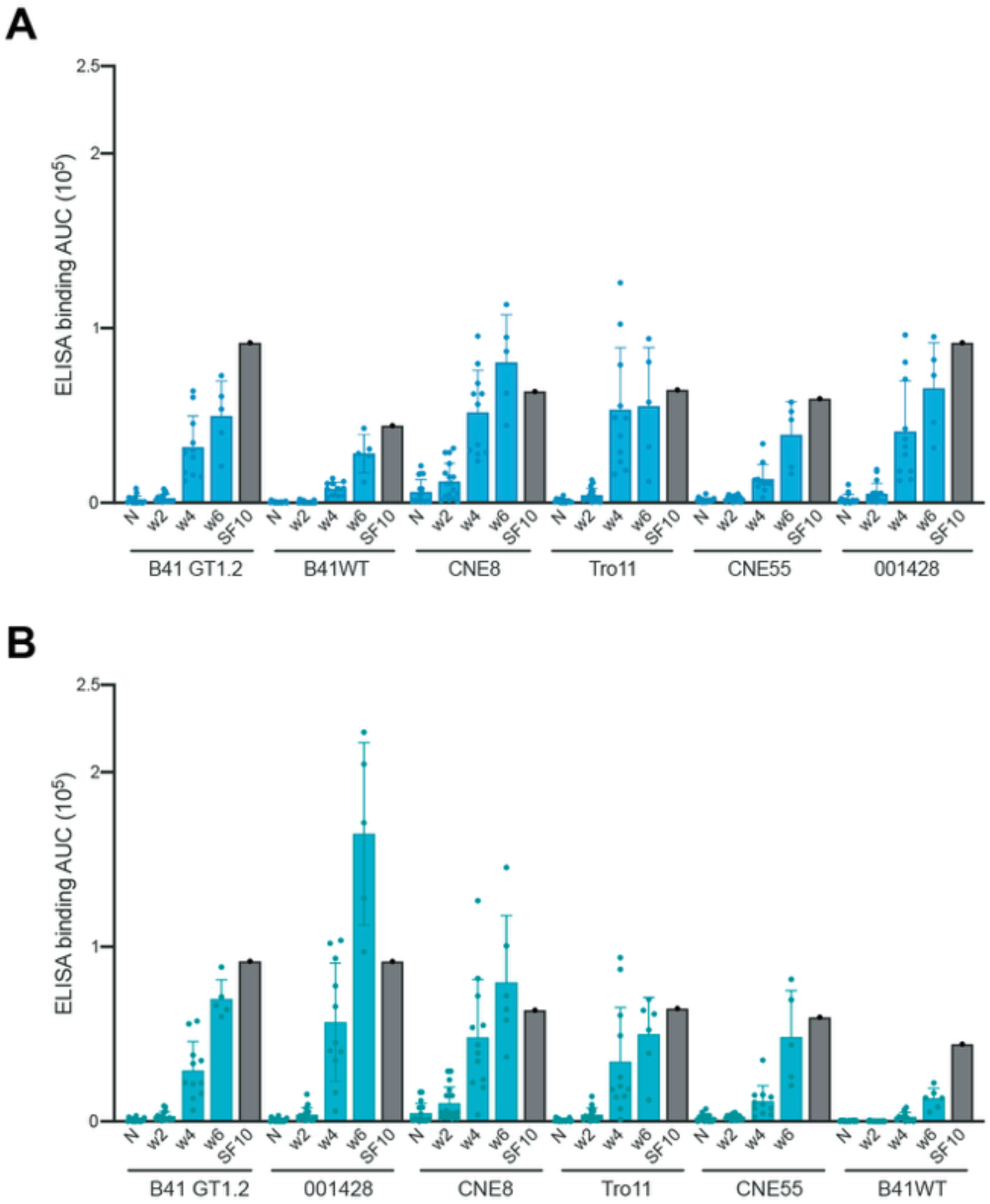
Longitudinal ELISA analysis of serum from the sequentially immunized SF10VHrev Ig KI mice. A,B) ELISA results showing binding of serum collected at w0 (naïve, N), two weeks after the prime (w2), 2 weeks after the first boost (w4) and 2 weeks after the second boost (w6) from mice in G1 (A) and G2 (B) against the immunogens used for each immunization and 3 heterologous HIV-1 env proteins. Black bars show binding of the SF10 bNAb. Each dot represents one mouse G1 n=17 G2 n=18.

**Supplementary Figure 6.**
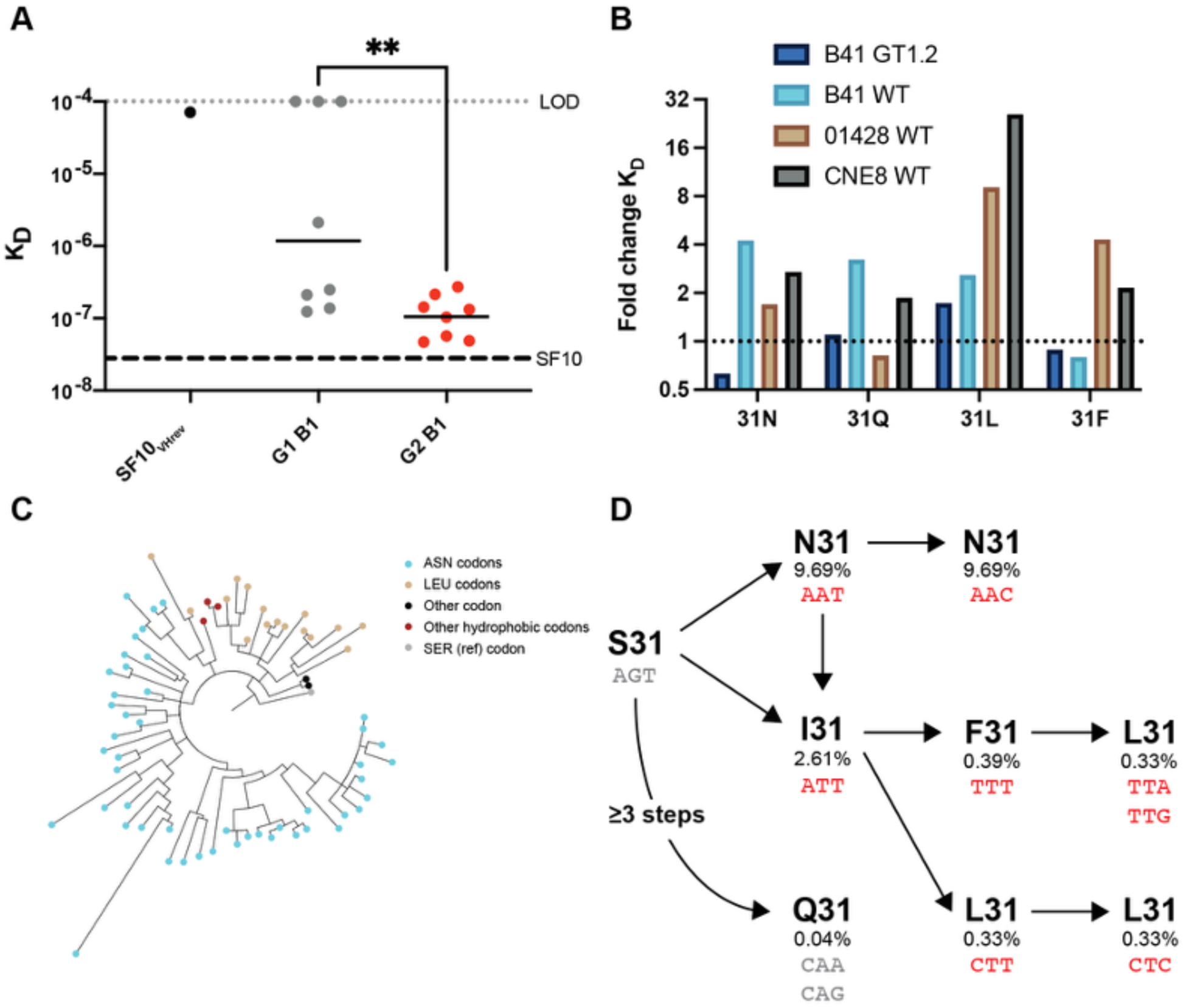
Affinity and codon accessibility influences evolutionary trajectories of SF-directed antibodies at position 31. **A)** K_D_ measured by BLI for mAb binding to CNE8 gp120c. Statistical testing was performed using Wilcoxon rank sum, ** p<0.01. Affinity of SF10 is shown as a dashed line for comparison. **B)** Fold change *k_off_* for reversion of somatic mutations at position 31 vs minSF10.11 background. **C)** Phylogenetic tree of sequences from mouse AJV8 (Group 2 Week 6) demonstrating separation of L31 and N31 containing branches of the tree. Tips are colored by amino acid at position 31 as shown. **D)** Potential paths to observed amino acid mutations and codons from the germline serine codon. Each arrow is a single base change unless otherwise indicated. Observed codons are highlighted in red. The percent chance of obtaining this codon in a similarly mutated antibody to SF10 (as determined by ARMADILLO) is noted. The high accessibility and affinity improvement for N31 toward B41 explains the strong enrichment observed in Figure 6. Similarly, the affinity enhancements for L31 and F31 with 01428 likely explain the enrichment of hydrophobic amino acids observed at week 4 in Group 2 mice.

**Table S1.**
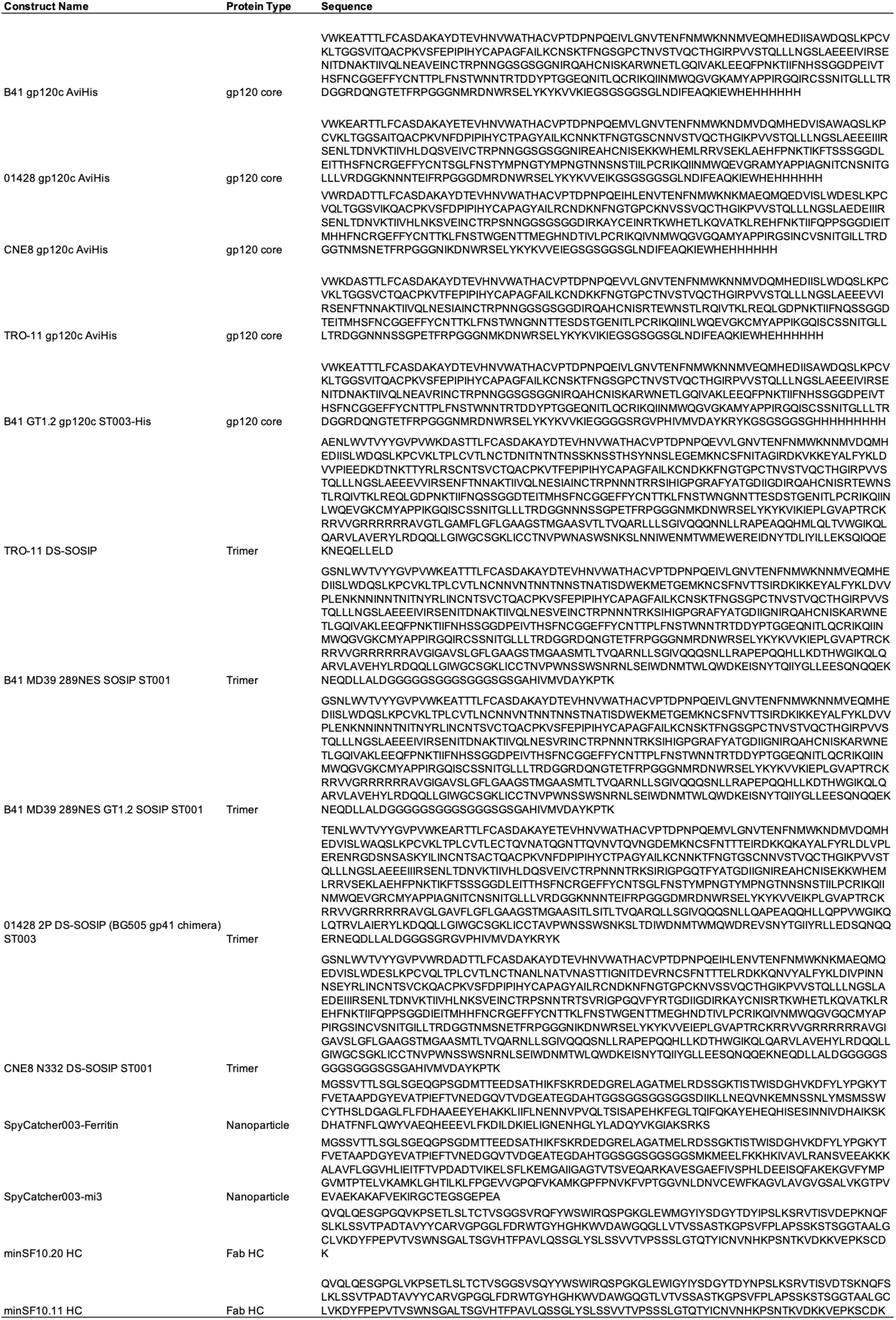
Protein sequences used in this study.

**Table S2.**
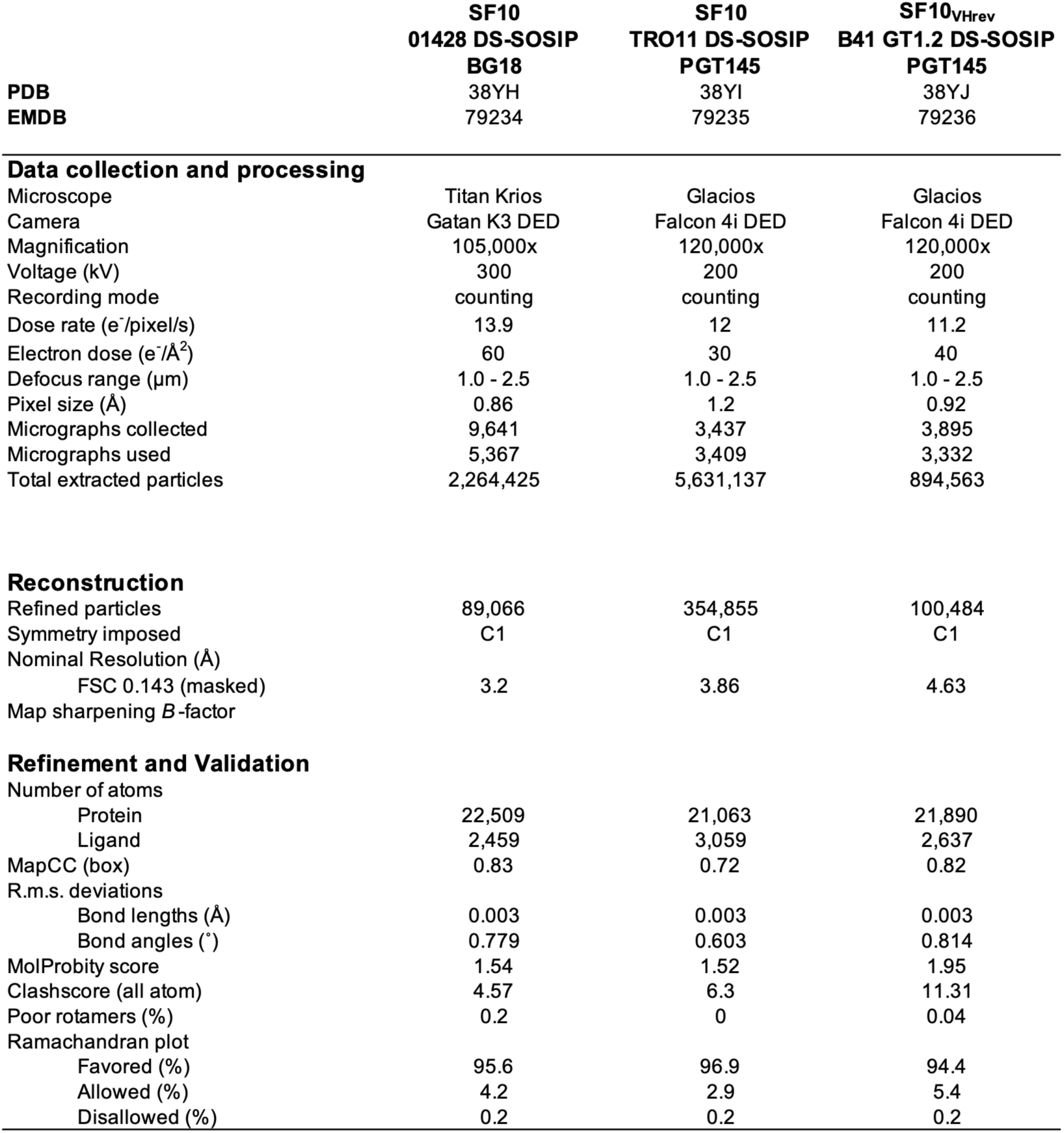
Cryo-EM data collection and refinement statistics.

**Table S3.**
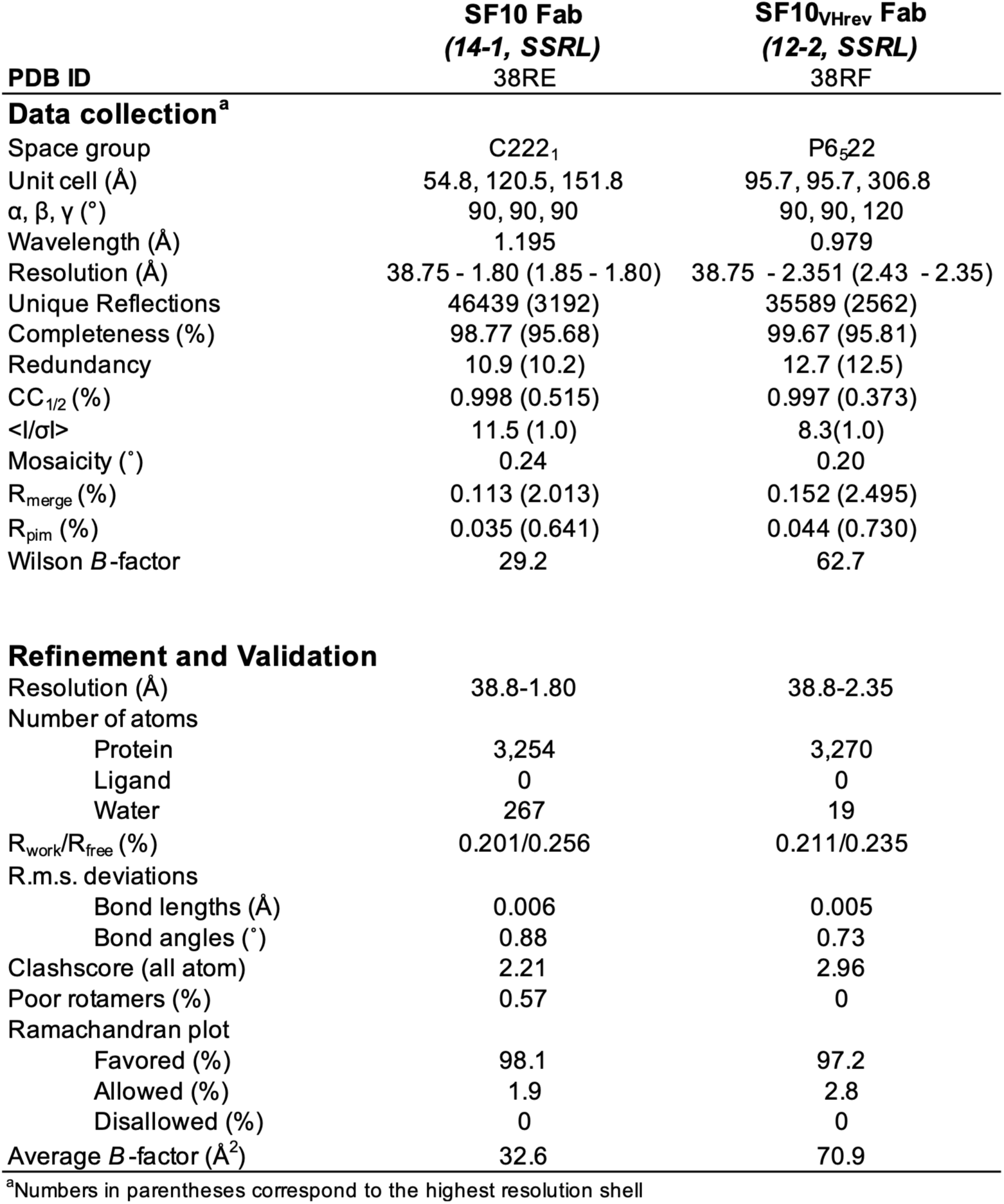
X-ray data collection and refinement statistics.

**Table S4.**
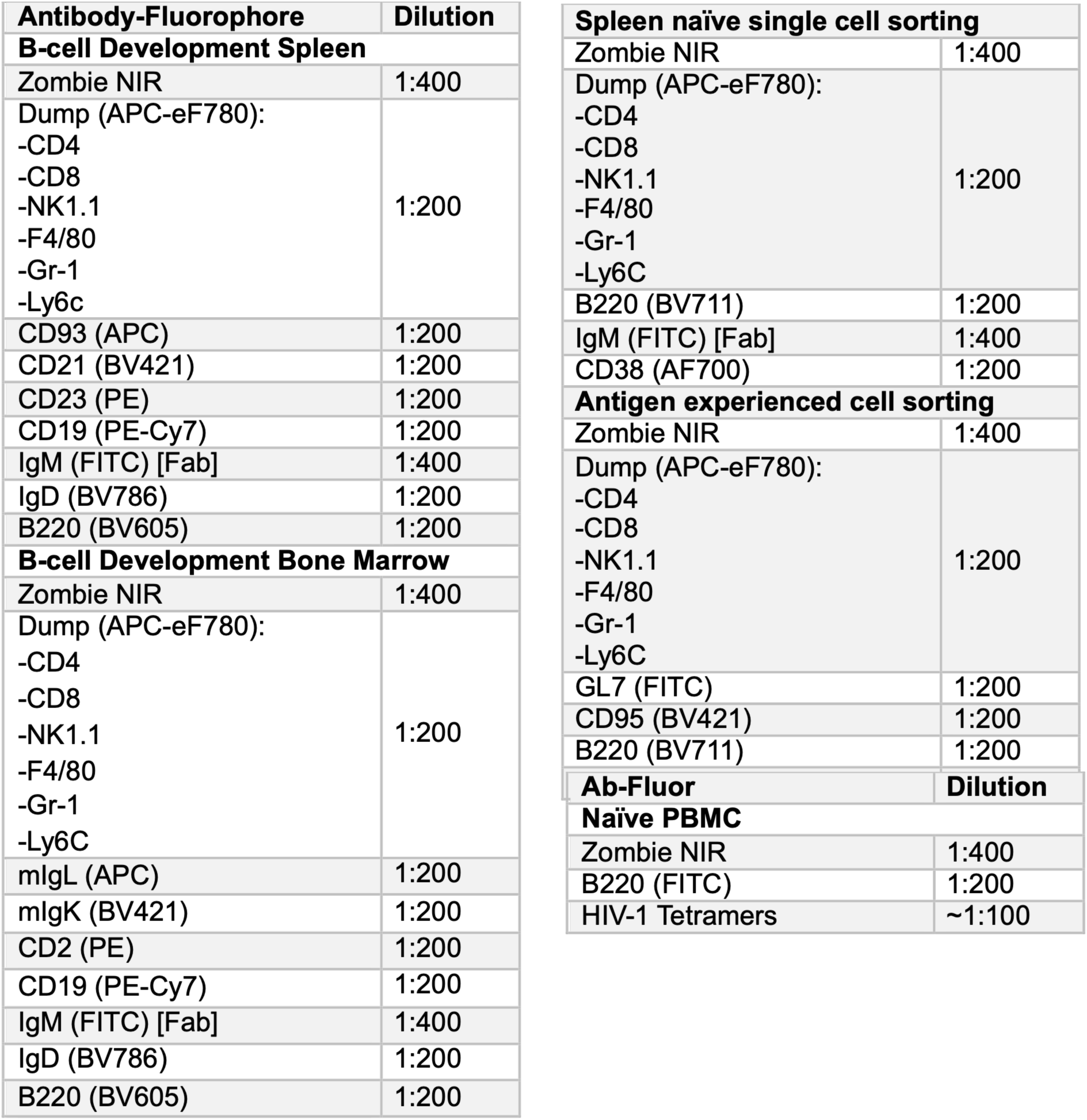
Antibody staining parameters.

**Table S5.**
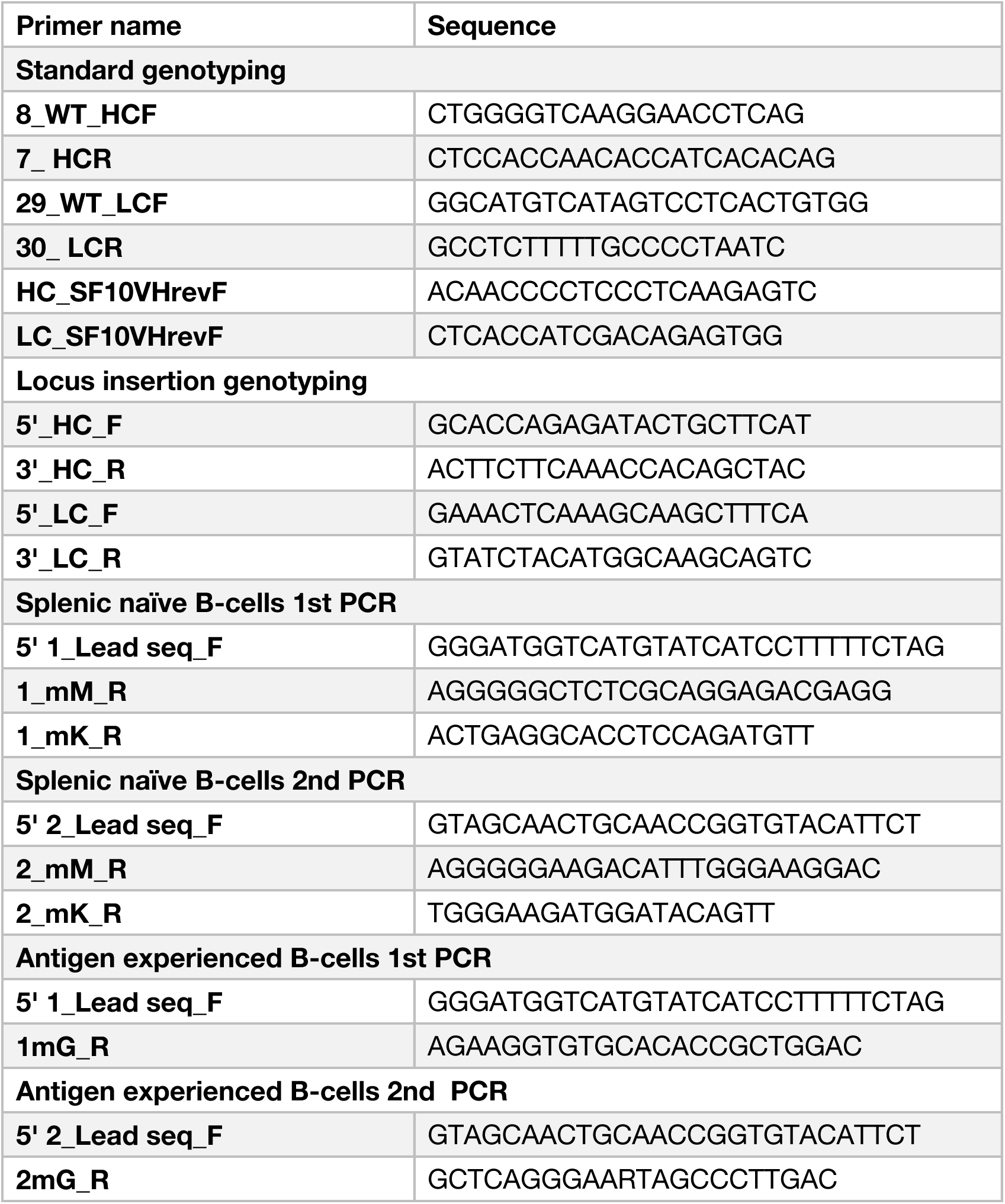
Primer sequences used in this study.

## REFERENCES

1. Transforming our world: the 2030 Agenda for Sustainable Development Preamble (2015)., 10.1057/978-1-137-45443-0_24 10.1057/978-1-137-45443-0_24.

2. Corey, L., and Gray, G.E. (2017). Preventing acquisition of HIV is the only path to an AIDS-free generation. 114, 3798–3800. 10.1073/pnas.1703236114.

3. Haynes, B.F., Wiehe, K., Borrow, P., Saunders, K.O., Korber, B., Wagh, K., McMichael, A.J., Kelsoe, G., Hahn, B.H., Alt, F., et al. (2023). Strategies for HIV-1 vaccines that induce broadly neutralizing antibodies. Nat. Rev. Immunol. 23. 10.1038/s41577-022-00753-w.

4. Wiehe, K., Saunders, K.O., Stalls, V., Cain, D.W., Venkatayogi, S., Martin Beem, J.S., Berry, M., Evangelous, T., Henderson, R., Hora, B., et al. (2024). Mutation-guided vaccine design: A process for developing boosting immunogens for HIV broadly neutralizing antibody induction. Cell Host Microbe 32, 693–709.e7. 10.1016/j.chom.2024.04.006.

5. Wagh, K., and Seaman, M.S. (2023). Divide and conquer: broadly neutralizing antibody combinations for improved HIV-1 viral coverage. Curr. Opin. HIV AIDS 18, 164–170. 10.1097/COH.0000000000000800.

6. Steichen, J.M., Lin, Y.C., Havenar-Daughton, C., Pecetta, S., Ozorowski, G., Willis, J.R., Toy, L., Sok, D., Liguori, A., Kratochvil, S., et al. (2019). A generalized HIV vaccine design strategy for priming of broadly neutralizing antibody responses. Science (80-. ). 366, 1–37. 10.1126/science.aax4380.

7. Escolano, A., Steichen, J.M., Dosenovic, P., Kulp, D.W., Golijanin, J., Sok, D., Freund, N.T., Gitlin, A.D., Oliveira, T., Araki, T., et al. (2016). Sequential Immunization Elicits Broadly Neutralizing Anti-HIV-1 Antibodies in Ig Knockin Mice. Cell 166, 1445–1458.e12. 10.1016/j.cell.2016.07.030.

8. Duan, H., Nkolola, J.P., Wang, S., Chaudhari, J., Teng, I.T., Lavine, C., Parchment, D.K., Sellers, G.S., McKee, K., O’Dell, S., et al. (2026). Vaccine elicitation of HIV-1 neutralizing antibodies against both V2 apex and fusion peptide in rhesus macaques. Cell Rep. 45, 116905. 10.1016/j.celrep.2025.116905.

9. Sutton, H.J., Ma, K.M., Steichen, J.M., Schiffner, T., Altheide, T.K., Liguori, A., Lu, D., Kubitz, M., Georgeson, E., Phelps, N., et al. (2025). Simultaneous Induction of Multiple Classes of Broadly Neutralizing Antibody Precursors by Combination Germline-Targeting Immunization in Non-Human Primates. Sci. Immunol. 10. 10.1126/sciimmunol.adu8878.Simultaneous.

10. Haynes, B.F., Saunders, K.O., Hahn, B.H., Wiehe, K., Baden, L.R., and Shaw, G.M. (2025). Status of HIV vaccine development: progress and promise. J. Int. AIDS Soc. 28, 4–6. 10.1002/jia2.26489.

11. Agudelo, M., Palus, M., Keeffe, J.R., Bianchini, F., Svoboda, P., Gazumyan, A., Cipolla, M., Kapoor, T., Guidetti, F., Yao, K., et al. (2021). Broad and potent neutralizing human antibodies to tick-borne flaviviruses protect mice from disease. 218.

12. Zhou, T., Zheng, A., Baxa, U., Chuang, G.Y., Georgiev, I.S., Kong, R., O’Dell, S., Shahzad-ul-Hussan, S., Shen, C.H., Tsybovsky, Y., et al. (2018). A Neutralizing Antibody Recognizing Primarily N-Linked Glycan Targets the Silent Face of the HIV Envelope. Immunity 48, 500–513.e6. 10.1016/j.immuni.2018.02.013.

13. van Schooten, J., Schorcht, A., Farokhi, E., Umotoy, J.C., Gao, H., van den Kerkhof, T.L.G.M., Dorning, J., Rijkhold Meesters, T.G., van der Woude, P., Burger, J.A., et al. (2022). Complementary antibody lineages achieve neutralization breadth in an HIV-1 infected elite neutralizer. PLoS Pathog. 18, 1–28. 10.1371/journal.ppat.1010945.

14. Sok, D., and Burton, D.R. (2018). Recent progress in broadly neutralizing antibodies to HIV. Nat. Immunol. 19, 1179–1188. 10.1038/s41590-018-0235-7.

15. Schoofs, T., Barnes, C.O., Suh-Toma, N., Golijanin, J., Schommers, P., Gruell, H., West, A.P., Bach, F., Lee, Y.E., Nogueira, L., et al. (2019). Broad and Potent Neutralizing Antibodies Recognize the Silent Face of the HIV Envelope. Immunity 50, 1513–1529.e9. 10.1016/j.immuni.2019.04.014.

16. Beem, J.S.M., Venkatayogi, S., Haynes, B.F., and Wiehe, K. (2023). ARMADiLLO : a web server for analyzing antibody m utation pr obabilities. 51, 51–56.

17. Batista, F.D., and Neuberger, M.S. (1998). Affinity dependence of the B cell response to antigen: A threshold, a ceiling, and the importance of off-rate. Immunity 8, 751–759. 10.1016/S1074-7613(00)80580-4.

18. Guenaga, J., Ádori, M., Bale, S., Phulera, S., Zygouras, I., Schleich, F.A., Castro Dopico, X., Agrawal, S., Ota, M., Wilson, R., et al. (2026). Vaccination generates broadly cross-neutralizing antibodies to the HIV Env apex. Nature 654, 777–785. 10.1038/s41586-026-10429-3.

19. Steichen, J.M., Madden, P.J., Flynn, C.T., Phulera, S., Shil, M., Kalyuzhniy, O., Liguori, A., Kifude, C., Sewall, L.M., Cottrell, C.A., et al. (2026). Vaccination elicits HIV broadly neutralizing antibodies in primates (Springer US) 10.1038/s41586-026-10837-5.

20. Marchitto, L., Wagh, K., Roark, R.S., Coleon, S., Li, H., Skelly, A.N., Hogarty, M.P., Habib, R., Ding, W., Ayyanathan, K., et al. (2026). Enhanced B cell priming induces broadly neutralizing HIV-1 apex antibodies (Springer US) 10.1038/s41586-026-10838-4.

21. Caniels, T.G., Prabhakaran, M., Ozorowski, G., MacPhee, K.J., Wu, W., van der Straten, K., Agrawal, S., Derking, R., Reiss, E.I.M.M., Millard, K., et al. (2025). Precise targeting of HIV broadly neutralizing antibody precursors in humans. Science (80-. ). 389. 10.1126/science.adv5572.

22. Jardine, J.G., Sok, D., Julien, J., Briney, B., Adachi, Y., Diwanji, D., Hsueh, J., Jones, M., Kalyuzhniy, O., Kubitz, M., et al. (2016). Minimally Mutated HIV-1 Broadly Neutralizing Antibodies to Guide Reductionist Vaccine Design. 1–33. 10.1371/journal.ppat.1005815.

23. Dam, K.M.A., Gristick, H.B., Li, Y.E., Yang, Z., Gnanapragasam, P.N.P., West, A.P., Seaman, M.S., and Bjorkman, P.J. (2025). Mapping essential somatic hypermutations in a CD4-binding site bNAb informs HIV-1 vaccine design. Cell Rep. 44. 10.1016/j.celrep.2025.115713.

24. Henderson, R., Zhou, Y., Stalls, V., Wiehe, K., Saunders, K.O., Wagh, K., Anasti, K., Barr, M., Parks, R., Alam, S.M., et al. (2023). Structural basis for breadth development in the HIV-1 V3-glycan targeting DH270 antibody clonal lineage. 1–17. 10.1038/s41467-023-38108-1.

25. Swanson, O., Rhodes, B., Wang, A., Xia, S.M., Parks, R., Chen, H., Sanzone, A., Cooper, M., Louder, M.K., Lin, B.C., et al. (2021). Rapid selection of HIV envelopes that bind to neutralizing antibody B cell lineage members with functional improbable mutations. Cell Rep. 36, 109561. 10.1016/j.celrep.2021.109561.

26. Swanson, O.M., Zhang, Q.E., Van Itallie, E., Tian, M., Brown, A.R., Harris, C., Kapingidza, A.B., Rhodes, B., Smith, L.M., Venkatayogi, S., et al. (2025). An engineered immunogen activates diverse HIV broadly neutralizing antibody precursors and promotes acquisition of improbable mutations. Sci. Transl. Med. 17, 1–13. 10.1126/scitranslmed.adr2218.

27. Saunders, K.O., Wiehe, K., Tian, M., Acharya, P., Bradley, T., Munir Alam, S., Go, E.P., Scearce, R., Sutherland, L., Henderson, R., et al. (2019). Targeted selection of HIV-specific antibody mutations by engineering B cell maturation. Science (80-. ). 366. 10.1126/science.aay7199.

28. Henderson, R., Anasti, K., Manne, K., Stalls, V., Saunders, C., Bililign, Y., Williams, A., Bubphamala, P., Montani, M., Kachhap, S., et al. (2024). Engineering immunogens that select for speci fi c mutations in HIV broadly neutralizing antibodies. Nat. Commun., 1–20. 10.1038/s41467-024-53120-9.

29. Swanson, O., Rhodes, B., Wang, A., Xia, S., Parks, R., Sanzone, A., Cooper, M., Louder, M.K., Lin, B.C., Doria, N.A., et al. (2021). Rapid selection of HIV envelopes that bind to neutralizing antibody B cell lineage members with functional improbable mutations. Cell Rep. 36. 10.1016/j.celrep.2021.109561.Rapid.

30. Habib, A.R., Roark, R.S., Li, H., Connell, A.J., Michael, P., Sowers, K.J., Ayyanathan, K., Plante, S.J., Bibollet-ruche, F., Rando, M., et al. (2025). Env-antibody coevolution identifies B cell priming as the principal bottleneck to HIV-1 V2 apex broadly neutralizing antibody development Affiliations :

31. Joyce, M.G., Georgiev, I.S., Yang, Y., Druz, A., Geng, H., Chuang, G.Y., Kwon, Y. Do, Pancera, M., Rawi, R., Sastry, M., et al. (2017). Soluble Prefusion Closed DS-SOSIP.664-Env Trimers of Diverse HIV-1 Strains. Cell Rep. 21, 2992–3002. 10.1016/j.celrep.2017.11.016.

32. Punjani, A., Rubinstein, J.L., Fleet, D.J., and Brubaker, M.A. (2017). cryoSPARC : algorithms for rapid unsupervised cryo-EM structure determination. 14. 10.1038/nmeth.4169.

33. Scheres, S.H.W. (2012). RELION : Implementation of a Bayesian approach to cryo-EM structure determination. J. Struct. Biol. 180, 519–530. 10.1016/j.jsb.2012.09.006.

34. Punjani, A., Zhang, H., and Fleet, D.J. (2020). Non-uniform refinement: adaptive regularization improves single-particle cryo-EM reconstruction. Nat. Methods 17. 10.1038/s41592-020-00990-8.

35. Levy, A., Raghu, R., Feathers, J.R., Grzadkowski, M., Poitevin, F., Johnston, J.D., Vallese, F., Clarke, O.B., Wetzstein, G., and Ellen, D. (2025). CryoDRGN-AI : Neural ab initio reconstruction of challenging cryo-EM and cryo-ET datasets. 1–36.

36. Jumper, J., Evans, R., Pritzel, A., Green, T., Figurnov, M., Ronneberger, O., Tunyasuvunakool, K., Bates, R., Žídek, A., Potapenko, A., et al. (2021). Highly accurate protein structure prediction with AlphaFold. Nature 596. 10.1038/s41586-021-03819-2.

37. Meng, E.C., Goddard, T.D., Pettersen, E.F., Pearson, Z.J., Morris, J.H., Ferrin, T.E., and Couch, G.S. (2023). UCSF ChimeraX : Tools for structure building and analysis. 1–13. 10.1002/pro.4792.

38. Adams, P.D. (2018). Real-space refinement in PHENIX for cryo-EM and crystallography research papers. 531–544. 10.1107/S2059798318006551.

39. Emsley, P., Lohkamp, B., Scott, W.G., and Cowtan, K. (2010). Features and development of Coot. Acta Crystallogr., 486–501. 10.1107/S0907444910007493.

40. Williams, C.J., Headd, J.J., Moriarty, N.W., Prisant, M.G., Videau, L.L., Deis, L.N., Verma, V., Keedy, D.A., Hintze, B.J., Chen, V.B., et al. (2017). MolProbity : More and better reference data for improved all-atom structure validation. 27, 293–315. 10.1002/pro.3330.

41. Wu, T. Te, and Kabat, E.A. (1970). AN ANALYSIS OF THE SEQUENCES OF THE VARIABLE REGIONS OF BENCE JONES PROTEINS AND MYELOMA LIGHT CHAINS AND THEIR IMPLICATIONS FOR ANTI-BODY COMPLEMENTARITY. J. Exp. Med. 132, 211–250.

42. Rubio, A.A., Baharani, V.A., Dadonaite, B., Parada, M., Abernathy, M.E., Wang, Z., Lee, Y.E., Eso, M.R., Phung, J., Ramos, I., et al. (2025). Bispecific antibodies targeting the N-terminal and receptor binding domains potently neutralize SARS-CoV-2 variants of concern. Sci. Transl. Med. 17. 10.1126/scitranslmed.adq5720.

43. Escolano, A., Gristick, H.B., Abernathy, M.E., Merkenschlager, J., Gautam, R., Oliveira, T.Y., Pai, J., Jr, A.P.W., Christopher, O., Cohen, A.A., et al. (2019). Immunization expands HIV-1 V3-glycan specific B-cells in mice and macaques. Nature 570, 468–473. 10.1038/s41586-019-1250-z.Immunization.

44. Krämer, S.D., Wöhrle, J., Rath, C., and Roth, G. (2019). Anabel : An Online Tool for the Real-Time Kinetic Analysis of Binding Events. 10.1177/1177932218821383.

45. Gristick, H.B., Hartweger, H., Loewe, M., van Schooten, J., Ramos, V., Oliveira, T.Y., Nishimura, Y., Koranda, N.S., Wall, A., Yao, K.H., et al. (2023). CD4 binding site immunogens elicit heterologous anti-HIV-1 neutralizing antibodies in transgenic and wild-type animals. Sci. Immunol. 8, eade6364. 10.1126/sciimmunol.ade6364.

46. Deiters, A. Bioconjugation Methods.

47. Wang, W., Liu, Z., Zhou, X., Guo, Z., Zhang, J., Zhu, P., Yao, S., and Zhu, M. (2019). Ferritin nanoparticle-based SpyTag/SpyCatcher-enabled click vaccine for tumor immunotherapy. Nanomedicine Nanotechnology, Biol. Med. 16, 69–78. 10.1016/j.nano.2018.11.009.

48. Sarzotti-kelsoe, M., Bailer, R.T., Turk, E., Lin, C., Bilska, M., Greene, K.M., Gao, H., Todd, C.A., Ozaki, D.A., Michael, S., et al. (2014). Optimization and Validation of the TZM-bl Assay for Standardized Assessments of Neutralizing Antibodies Against HIV-1. J. Immunol. Methods, 131–146. 10.1016/j.jim.2013.11.022.Optimization.

